# Unbiased construction of a temporally consistent morphological atlas of neonatal brain development

**DOI:** 10.1101/251512

**Authors:** Andreas Schuh, Antonios Makropoulos, Emma C. Robinson, Lucilio Cordero-Grande, Emer Hughes, Jana Hutter, Anthony N. Price, Maria Murgasova, Rui Pedro A. G. Teixeira, Nora Tusor, Johannes K. Steinweg, Suresh Victor, Mary A. Rutherford, Joseph V. Hajnal, A. David Edwards, Daniel Rueckert

**Affiliations:** Biomedical Image Analysis Group, Department of Computing, Imperial College London, London, United Kingdom; Centre for the Developing Brain, King’s College London, London, United Kingdom; Department of Biomedical Engineering, School of Biomedical Engineering and Imaging Sciences, King’s College London, London, United Kingdom

**Keywords:** dHCP, spatio-temporal, neonatal MRI, brain atlas

## Abstract

Premature birth increases the risk of developing neurocognitive and neurobehavioural disorders. The mechanisms of altered brain development causing these disorders are yet unknown. Studying the morphology and function of the brain during maturation provides us not only with a better understanding of normal development, but may help us to identify causes of abnormal development and their consequences. A particular difficulty is to distinguish abnormal patterns of neurodevelopment from normal variation. The Developing Human Connectome Project (dHCP) seeks to create a detailed four-dimensional (4D) connectome of early life. This connectome may provide insights into normal as well as abnormal patterns of brain development. As part of this project, more than a thousand healthy fetal and neonatal brains will be scanned *in vivo.* This requires computational methods which scale well to larger data sets. We propose a novel groupwise method for the construction of a spatio-temporal model of mean morphology from cross-sectional brain scans at different gestational ages. This model scales linearly with the number of images and thus improves upon methods used to build existing public neonatal atlases, which derive correspondence between all pairs of images. By jointly estimating mean shape and longitudinal change, the atlas created with our method overcomes temporal inconsistencies, which are encountered when mean shape and intensity images are constructed separately for each time point. Using this approach, we have constructed a spatio-temporal atlas from 275 healthy neonates between 35 and 44 weeks post-menstrual age (PMA). The resulting atlas qualitatively preserves cortical details significantly better than publicly available atlases. This is moreover confirmed by a number of quantitative measures of the quality of the spatial normalisation and sharpness of the resulting template brain images.

## 1. Introduction

Medical image analysis has made significant progress over the past decades. With advances in acquiring high quality images of the developing human brain using magnetic resonance imaging (MRI), analysing this data to understand brain development is rapidly becoming feasible (Kwon et al., 2014).

Many studies have shown that premature birth increases the risk of developing neurocognitive and neurobehavioural disorders (Rutherford, 2002; Blencowe et al., 2012; Sled and Nossin-Manor, 2013). A known difference to term-born infants is the reduced cortical folding observed in premature-born infants at term-equivalent age (Huppi et al., 1996; Inder et al., 2005; Kapellou et al., 2006). Other reported abnormalities include: (1) white matter (WM) signal abnormalities such as lesions and diffuse excessive high signal intensities, (2) cortical grey matter (cGM) signal abnormalities, and (3) cerebellar hemorrhages (cf. Kwon et al., 2014, and references therein). However, the mechanisms of these alterations are unknown (Molnar and Rutherford, 2013). Studying the morphology and function of the brain during maturation, provides us not only with a better understanding of normal development, but may help identify causes for these factors.

The Developing Human Connectome Project (dHCP) seeks to create a 4D connectome of early life. Understanding this connectome may provide insights into abnormal patterns of brain development. In this context, an atlas of normal brain development is important, as it facilitates spatial normalisation, voxel-based morphometry (Ashburner and Friston, 2000), deformation-based morphometry (Ashburner et al., 1998), and computational anatomy studies (Grenander and Miller, 1998; Thompson and Toga, 2002).

In previous work (Schuh et al., 2014), we constructed a spatio-temporal neonatal atlas from registrations between all pairs of images, and the Log-Euclidean mean of stationary velocity fields (SVFs) (Arsigny et al., 2006). It improved upon the method of Serag et al. (2012a), which is based on the freeform deformation (FFD) algorithm (Rueckert et al., 1999), and arithmetic mean of displacement fields. A longitudinal model of brain morphology is hereby obtained using Nadaraya-Watson kernel regression similar to Davis et al. (2007), with adaptive kernel size as proposed by Serag et al. (2012a).

A drawback of these techniques is the quadratic computational complexity with respect to the number of images, which means that these approaches are difficult to scale to large datasets. A more significant limitation is the reliance on accurate pairwise correspondences, and the dependence on average spatial correspondences, which can capture complex cortical folding patterns. The less accurate the individual registrations are, the less detail is preserved. An approach which iteratively refines the transformations that relate each anatomy with the average space, may be more robust towards residual misalignment.

Therefore, instead of expressing the spatio-temporal atlas in terms of the transformations relating each brain image to all others, in this work we derive a new recursive and continuous formulation based on temporal kernel regression, the transformations relating each image to the spatio-temporal atlas space, and their respective age-dependent Log-Euclidean mean. This novel approach jointly estimates mean shape and longitudinal change iteratively. A comparison of our methods to the atlas obtained with Serag et al.’s (2012a) technique demonstrates a marked increase in anatomical detail.

### 1.1 Overview

In section 1.2, we review main approaches to brain atlas construction. In section 1.3, we summarise related work in which an average model of the neonatal brain has been created. Our method for the construction of a spatio-temporal atlas of neonatal brain morphology is outlined in section 2. Here, we first detail the initial affine spatial normalisation of all brain images in section 2.2. The unbiased diffeomorphic registration method, which is the basis for our deformable atlas construction, is described in section 2.5. The atlas construction itself is derived in section 2.6. We compare our method to previous approaches in section 3, and discuss these results in section 4.

### 1.2 Brain atlas construction

Figure 1a illustrates an atlas construction referred to herein as “pairwise” technique. First, the transformations between all pairs of images (squares) are computed (blue lines). A mean transformation (green arrows) is then derived for each image, which maps it to the average space (circle). Different approaches distinguish themselves by the transformation model, registration method used, and the definition of a mean of pairwise correspondences.

**Figure 1.**
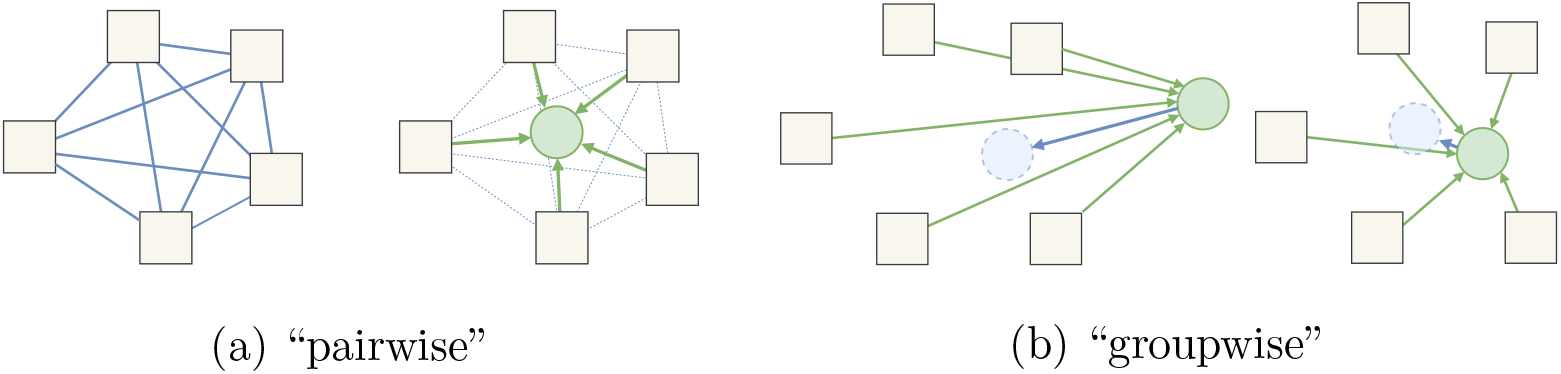
Illustration of common approaches to single-time point brain atlas construction. A spatio-temporal model is obtained using kernel regression along the time dimension.

An adult brain atlas using a pairwise method with affine transformations was first constructed by Woods et al. (1998). A non-linear extension was proposed by Seghers et al. (2004) based on the arithmetic mean of displacement fields obtained by non-parametric deformable registration. Similarly, the neonatal atlas construction of Serag et al. (2012a) employs the arithmetic mean of displacement-based FFDs, which are neither inverse consistent nor guaranteed diffeomorphic. These limitations were overcome by our previous pairwise approach (Schuh et al., 2014), which is based on an efficient diffeomorphic registration method, and the Log-Euclidean mean of SVFs.

Two steps of a “groupwise” approach, on the other hand, are depicted in figure 1b (left to right). At each iteration, the individual anatomies (squares) are registered to a reference anatomy (green circle). The average image is subsequently deformed by the average of the deformations (blue arrow) to form a new mean brain shape and intensity model (dashed blue circle), which is the reference for the next iteration (right). This iteratively minimises the distance of the reference to each anatomy until the residual deformation approaches zero. Such atlas construction resembles groupwise registration techniques (e.g. Studholme, 2003; Studholme and Cardenas, 2004; Bhatia et al., 2004, 2007; Zollei, 2006; Noblet et al., 2008; Wu et al., 2012).

A groupwise atlas construction, where one of the brain anatomies is used as initial reference, was first proposed by Guimond et al. (1998). Unbiased methods based on large deformation models similar to large deformation diffeomorphic metric mapping (LDDMM) (Beg et al., 2005) were pioneered by Joshi et al. (2004), Lorenzen et al. (2005, 2006), and Avants et al. (2010). Of these, the initial greedy approach of Joshi et al. (2004) and its non-greedy variant of Lorenzen et al. (2005) are based on the sum of squared differences, and therefore unsuitable for the construction of an atlas of the developing brain due to wide intensity variations. The symmetric groupwise normalisation (SyGN) of Avants et al. (2010), on the other hand, is based on the symmetric normalisation (SyN) method (Avants et al., 2008), which uses local normalised cross-correlation (LNCC) as image dissimilarity measure, so it may be more directly applicable to brain images of the developing brain.

### 1.3 Neonatal brain atlases

Kazemi et al. (2007) created the first population average from seven neonatal brains scanned at 39 to 42 weeks PMA for spatial normalisation using the Statistical Parameter Mapping (SPM) toolbox (Ashburner and Friston, 1999). Validation of spatial normalisation using this template instead of the adult template created at the Montreal Neurological Institute (MNI) on another seven brain scans showed that a neonatal template results in better co-registration. The same research group also provided a symmetric template built from all 14 term-born neonates for asymmetry studies of language development (Noorizadeh et al., 2013). Recently, a bimodal neonatal brain template consisting of an average magnetic resonance (MR) and computed tomography (CT) image was added (Ghadimi et al., 2017).

A set of 14 T1-weighted (T1w), 20 T2-weighted (T2w), and 20 diffusion tensor imaging (DTI) scans acquired within 38 to 41 weeks PMA were used by Oishi et al. (2011) to create different templates after affine and nonrigid alignment. Non-rigid registration was done using dual-channel LDDMM (Ceritoglu et al., 2009) to match the mean diffusivity (MD) and fractional anisotropy (FA) maps derived from the DTI scans.

A parcellation of the cortex of 10 healthy neonates born at term, and scanned between 40 and 43 weeks PMA, which is consistent with the cortical and subcortical adult brain atlas used in FreeSurfer (Desikan et al., 2006), was done by Alexander et al. (2016). Structural T1w and T2w templates for spatial normalisation to this cortical parcellation atlas were created using SyGN (Avants et al., 2010) of the Advanced Normalization Tools (ANTs).

Kuklisova-Murgasova et al. (2011) created an affine spatio-temporal atlas consisting of T2w average images of 142 preterm-born neonates and corresponding tissue probability maps for 29 to 44 weeks PMA. Using the non-rigid pairwise approach, Serag et al. (2012b) built a spatio-temporal model of average T1w and T2w intensities from 204 preterm-born neonates, also with accompanying tissue probability maps. Makropoulos et al. (2016) furthermore non-rigidly registered automatic whole brain segmentations of 87 brain structures of 420 mostly preterm-born neonates to the age-matched T2w templates created by Serag et al. (2012b), in order to build a fine-granular probabilistic atlas. These automatic segmentations are based on 20 single-subject atlases of 15 preterm-born neonates scanned at term-equivalent age, and 5 term-born neonates, referred to as ALBERTs. Each image was manually segmented as described in Gousias et al. (2012) to create an atlas of 50 anatomical regions. These were further subdivided into cortical and subcortical parts by Makropoulos et al. (2016).

Table 1 summarises publications which created a neonatal brain atlas. Of these, only Kuklisova-Murgasova et al. (2011), Serag et al. (2012a,b), and Schuh et al. (2014) constructed a 4D model of mean shape and intensity. Atlases of the developing brain were also reviewed by Benkarim et al. (2017).

**Table 1.** Atlases of the neonatal human brain. Columns from left to right: Citation, imaging modalities (mod.), number of subjects (no), post-menstrual age in weeks (age), method used for temporal regression (temp.), and spatial normalisation approach. Acronyms: KR: kernel regression with constant kernel width; AKR: adaptive kernel regression; HAMMER: Hierarchical Attribute Matching Mechanism for Elastic Registration (Shen and Davatzikos, 2002); IRTK: Image Registration ToolKit (Rueckert et al., 1999).

|  | mod. | no | age | temp. | registration |
| --- | --- | --- | --- | --- | --- |
| Kazemi et al. (2007) | T1w | 7 | 39–42 | – | non-rigid, SPM2 |
| Noorizadeh et al. (2013) | T1w | 14 | 39–42 | – | non-rigid, SPM8 |
| Ghadimi et al. (2017) | T1w | 7 | 39–42 | – | non-rigid, SPM8 |
|  | CT |  |  |  |  |
| Oishi et al. (2011) | T1w | 14 | 38–41 | – | dual-channel |
|  | T2w | 20 |  |  | LDDMM |
|  | DTI |  |  |  |  |
| Alexander et al. (2016) | T1w | 10 | 40–43 | – | SyGN, ANTs, |
|  | T2w |  |  |  | groupwise |
| Gousias et al. (2012) | T2w | 15 | 26–35 | – | – |
| Gousias et al. (2012) | T2w | 5 | 39–45 | – | – |
| Shi et al. (2011) | T1w | 95 | 39–46 | – | HAMMER, |
|  | T2w |  |  |  | groupwise |
| Kuklisova-Murgasova et al. (2011) | T2w | 142 | 29–44 | KR | affine, IRTK |
| Serag et al. (2012b) | T1w | 204 | 28–44 | AKR | FFD, IRTK, |
|  | T2w |  |  |  | pairwise |
| Schuh et al. (2014) | T2w | 118 | 28–44 | AKR | SVF, pairwise |
| Makropoulos et al. (2016) | T2w | 420 | 28–44 | AKR | FFD, IRTK |

## 2 Material and methods

### 2.1 Data acquisition

The image data was collected at St. Thomas Hospital, London, on a Philips 3T scanner using a dedicated 32 channel neonatal head coil (Hughes et al., 2017). T2w images were obtained using a turbo spin echo (TSE) sequence, acquired in two stacks of 2D slices in sagittal and axial planes, using parameters: TR = 12 s, TE = 156 ms, SENSE factor 2.11 (axial) and 2.58 (sagittal). Overlapping slices with resolution 0.8 × 0.8 × 1.6 mm^3^ were acquired. During motion corrected reconstruction (Cordero-Grande et al., 2017), the images are resampled to an isotropic voxel size of 0.5 mm. T1w images were acquired using an inversion recovery TSE sequence at the same resolutions with TR = 4.8 s, TE = 8.7 ms, SENSE factor 2.26 (axial) and 2.66 (sagittal). Babies were imaged without sedation during natural sleep. All images were checked for abnormalities by a paediatric neuroradiologist.

### 2.2 Image preprocessing

The reconstructed volumes were preprocessed using a pipeline designed for neonatal brain images, including an initial rough brain extraction (Smith, 2002) and bias field correction (Tustison et al., 2010). The bias corrected images are segmented into 87 regions of interest (ROIs) using an extension of the developing region annotation with expectation maximisation (Draw-EM) algorithm developed by Makropoulos et al. (2014). For details, including references to the software, please refer to Makropoulos et al. (2017).

### 2.3 Image selection

Out of 495 processed brain scans of 474 participants, we selected images of 275 neonates. Their age distributions are shown in figure 2. This set of images was obtained via a process of exclusion based on (a) manual image quality score between 1 (poor) and 4 (good) (Makropoulos et al., 2017), excluding images with score 1 or 2, (b) availability of both T1w and T2w images, (c) exclusion of follow up scans, (d) maximum time to scan after birth of 4 weeks, and (e) minimum age at scan of 35 weeks PMA.

**Figure 2.**
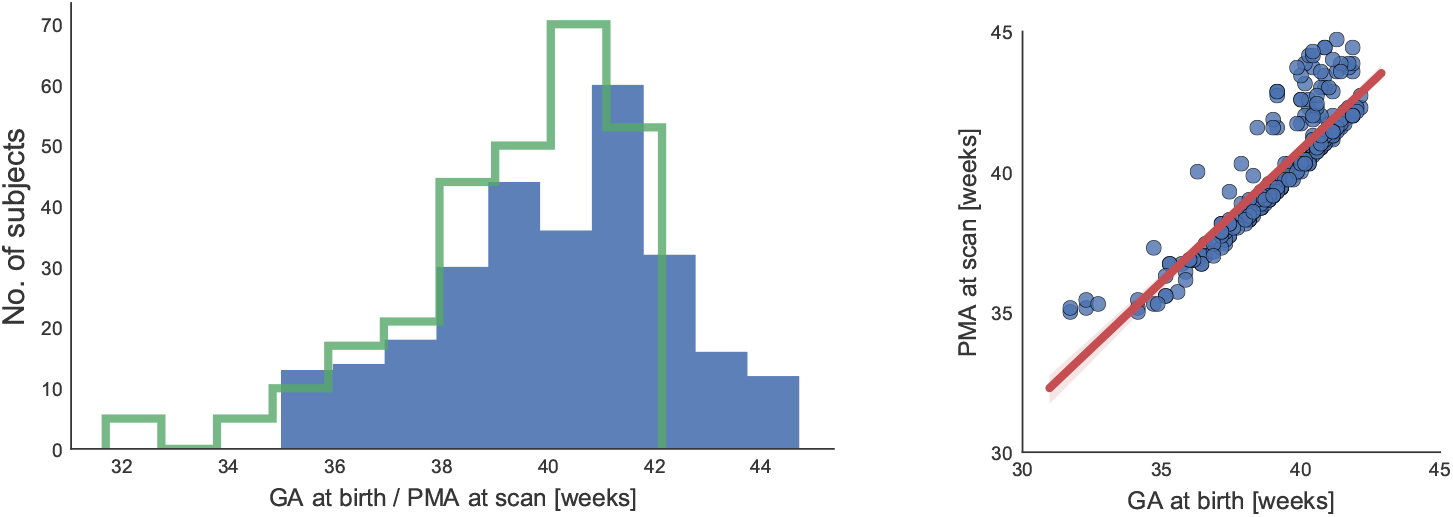
The histogram plot on the left shows the distribution of gestational age (GA) at birth (steps) and PMA at scan (bars) of the used dataset. The scatter plot on the right is overlayed with the regression line mapping from GA at birth to PMA at scan. Most neonates were scanned shortly after birth, with an average time to scan of 0.8 weeks.

### 2.4 Unbiased global normalisation

Pose differences of the imaged anatomies are removed by a rigid alignment to the 40 weeks template of the atlas constructed by Serag et al. (2012a). Differences in brain size are also removed at this global normalisation stage, such that the template construction based on deformable registration can focus on local shape differences. The so constructed template brain images are rescaled afterwards to reflect global volume changes.

The global normalisation and age-dependent rescaling is similar to the atlas construction proposed by Kuklisova-Murgasova et al. (2011), where only affine registration was employed to create average brain images and tissue probability maps of the preterm-born neonate. For this, the authors use a common reference image for the affine registrations. Bias towards the reference anatomy is hereby accounted for by composing each affine transformation with the inverse of their Log-Euclidean mean (Arsigny et al., 2007). Under the assumption of ideal, and thereby inverse-consistent and transitive affine transformations, the Log-Euclidean mean is independent of the chosen reference (Pennec and Arsigny, 2013). Bias towards the reference is thus removed by mapping it to the barycentre of all observed anatomies.

In practice, however, affine registration is more accurate for anatomies that are more similar to the reference anatomy. The choice of which of the two images is to remain fixed, and which is being transformed and resampled, further affects a pairwise registration (c.f. Cachier and Rey, 2000). These biases result in inconsistent transformations that are neither inverse-consistent nor transitive (Woods et al., 1998). Consequently, some anatomical bias is retained when a reference image is used for affine pre-alignment.

To eliminate this remaining bias, we perform affine registrations between all pairs of images, i.e., select each image in turn as reference. A mean transformation diferent from the Log-Euclidean mean employed in our work, which is derived from affine transformations between all pairs of images, was first employed by Woods et al. (1998) to create an adult brain template. Due to lack of inverse-consistency, some residual global misalignment remains, however. We therefore repeat the steps detailed below for another iteration.

Note that, while it would have been possible to refine through less costly, direct registrations of all images to the initial average, the benefits of a robust and unbiased pre-alignment of the images, which in turn impacts the subsequent deformable registrations, outweigh the added computational cost.

Formally, the *k*-th homogeneous coordinate transformation, which maps image *I_i_* to the unbiased global reference space, is given by the recursion

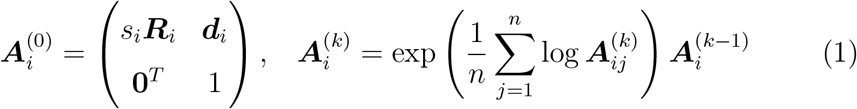

where 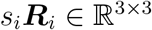 is a rotation followed by isotropic scaling, and 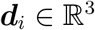 is a translation. These are found by initial rigid alignment of the i-th image to a selected reference image. The pairwise transformations 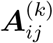are found by affine registration, as detailed in Appendix B. The matrix exponential and logarithm are denoted by exp and log (Alexa, 2002; Arsigny et al., 2007).

After *K* iterations,

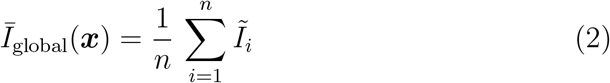

is the unbiased population-specific average image, where 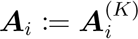, and

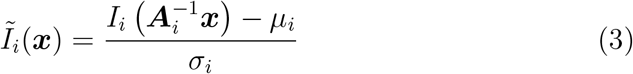

are the brain images after global spatial normalisation, with zero mean intensity and unit variance, whereby the original mean intensity and standard deviation are denoted by *μ_i_* and *σ_i_*, respectively. The normalisation of intensities by z-score compensates for global differences in intensity distributions.

### 2.5 Diffeomorphic image registration

The atlas construction is based on a fast and unbiased stationary velocity free-form deformation (SVFFD) algorithm detailed in the following.

#### 2.5.1 Parametric deformation model

The diffeomorphic mapping between a pair of images is given by the group exponential of a SVF, which in our work is a parametric spline function ***υ(x)*** as defined in Appendix A. A spatial mapping 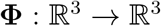 is then given by the group exponential, i.e., Φ = exp (*υ*), which is the solution at time *t* = 1 of the stationary ordinary differential equation (ODE) (Arsigny et al., 2006)

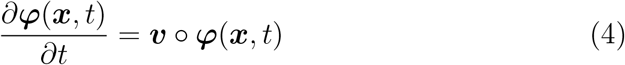

with initial condition *φ*(*x*, 0) = *x*. Hence, a SVF is the infinitesimal generator of a one-parameter subgroup of diffeomorphisms *φ_t_* = exp (*t*·*υ*).

The cubic B-spline parametrisation of the SVF was first proposed for brain image registration by Modat et al. (2012). It inherits some of the benefits of the not-guaranteed-diffeomorphic FFD model (Sederberg and Parry, 1986; Rueckert et al., 1999), namely an implicit smooth interpolation of the velocity field, and a parametric representation, which reduces the number of parameters and allows for an analytic derivation. Note that even though the support of the basis functions is limited, a control point may still affect the deformation of a point outside its local support region. This can be explained by considering the trajectory of a particle subjected to the flow induced by the SVF. Even when this trajectory originates at a position outside the local support region of a control point, it may enter it at a later time point *t* > 0.

The flow *φ_t_*, and therefore the group exponential, can be computed using a numerical integration scheme. For example, Pai et al. (2015) use a forward Euler integration. The advantage of a SVF over a time-varying velocity field is due to a method known as scaling and squaring (SS), i.e.,

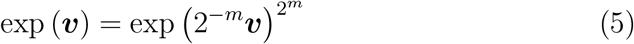

where exp (*υ*)^2^ = exp (*υ*) o exp (*υ*) and exp (2^*−m*^*υ*) ≈ Id +2^*−m*^*υ* (Arsigny et al., 2006): Instead of *m* Euler steps, only log_2_ *m* squaring steps are needed. To further reduce the computational cost of the integration, a SS on the coarser control point lattice of *υ* was proposed by Modat et al. (2012). Different methods for computing exp(*υ*) were compared by Bossa et al. (2008).

In Modat et al. (2012), it is noted that the SVF must be sufficiently smooth for the exponential to be a diffeomorphic mapping. Regularisation terms based on the Jacobian determinant of the SVF are therefore used. Given the cubic B-spline parametrisation, we can always find a minimum number of squaring steps, which guarantees positive Jacobian determinants of the scaled velocity field. This is because the Jacobian determinant of Φ at *x* is the product of determinants along the trajectory of a particle originating at this point. By choosing the integration step such that the maximum scaled coefficient is less than 0.4 times the minimum control point spacing, all factors are positive, and a diffeomorphic mapping is ensured (c.f. Choi and Lee, 2000; Rueckert et al., 2006). Folding and inconsistencies between exp(*υ*) and exp(–*υ*) may still be observed due to the numerical integration, i.e., the errors introduced during the composition of discrete deformation fields at each squaring step. Numerical errors caused by the integration are reduced by avoiding extreme Jacobian determinant values (Modat et al., 2012).

#### 2.5.2 Symmetrie energy function

Having defined the SVFFD model, we now detail our unbiased pairwise image registration method based on a symmetric energy formulation. Unlike the closely related method of Modat et al. (2012), where forward and backward mappings are parametrised by two separate SVFFDs, our formulation is based on a single spline function. Similar to the consistent-midpoint-cost proposed for non-stationary LDDMM by Beg and Khan (2007), we define

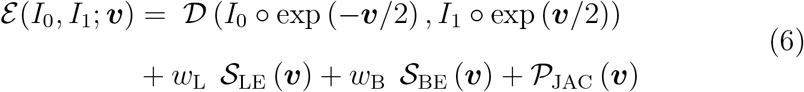

where *I*_{0;1}_ are the images to be registered. For neonatal brain image registration, suitable choices of image dissimilarity measures *D* are normalised mutual information (NMI), and LNCC. In case of LNCC, our symmetric data term is equivalent to the data matching term of the non-parametric LCC LogDemons approach (Lorenzi et al., 2013). To explicitly enforce smoothness of the SVF, regularisation terms based on first and second order derivatives are employed. These are the elastic potential of the spline-based SVF (similar to the non-parametric viscous-fluid model (Christensen et al., 1996)), i.e.,

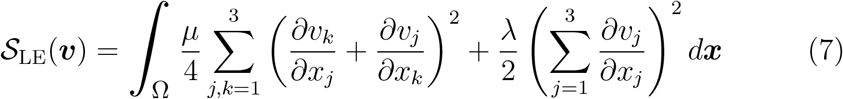

and the bending energy used in the FFD algorithm (Rueckert et al., 1999), but applied here to the SVF instead of a displacement field, i.e.,

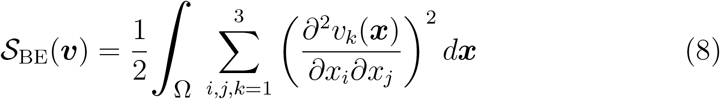

The non-negative weights *w_L_* and *w_B_* determine the trade-off between the smoothness imposed on the SVF and the image matching term. In addition to the explicit smoothness constraints, penalty terms based on the Jacobian determinants of the SVF can be added. A weighted sum of Jacobian-based penalty terms is denoted in equation (6) by 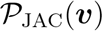. It should be noted that the bending energy and linear elasticity constraints are symmetric due to the squaring of derivatives, i.e., 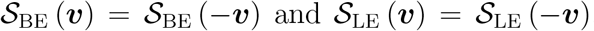. Jacobian-based penalties are generally not symmetric, however, and must be evaluated for both the positive and negative SVF. Therefore,

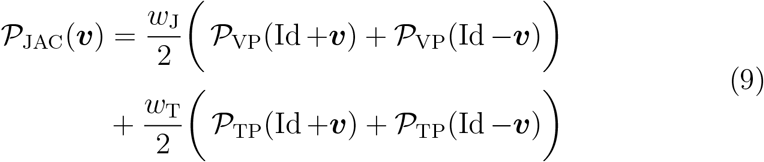

where 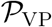and 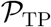are penalties of the deformation induced by the instantaneous velocity field and its approximate inverse, respectively, i.e.,

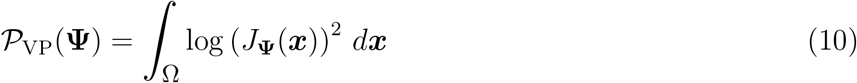

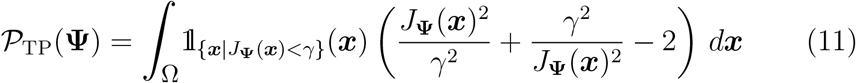

where the Jacobian determinant of a mapping Ψ is denoted by *J*_Ψ_(*x*), and the indicator function 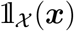is one for *x* ∊ *X*, and zero otherwise. While 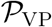penalises both excessive contraction and expansion, 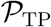is active for determinants *J* < γ < 1. It was used by Rueckert et al. (2006) as topology-preservation constraint in case of the FFD algorithm. Note that 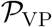is undefined for *J* ≤ 0. Similarly, 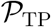is undefined for *J* = 0, and has the undesired property that it can prevent already negative determinants from being corrected for, since it is monotone increasing for *J* ∊ [−γ, 0).

Different implementations may handle the case of non-positive Jacobian determinants differently. An approach found in the Image Registration ToolKit (IRTK)^1^ is to simply clamp the values below a threshold ∊, e.g., ∊ = 10^−6^. This has the disadvantage that the value of the energy function does not change as long as the Jacobian determinant is less than ∊. This may result in a premature termination of the iterative optimisation, because the total energy may not have improved when stepping along the gradient direction. A different strategy to avoid undefined values of 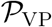 is taken by NiftyReg. After every gradient step, non-positive Jacobian determinants are detected, and an iterative unfolding, which aims to remove these based on a separate maximisation of their values, is performed (Modat et al., 2012). This strategy may fail to resolve all non-positive values, which then results in undefined penalties and may in consequence prematurely terminate the registration.

We instead define both Jacobian-based penalty terms as piecewise functions, where the functions above a small positive threshold *∊* are identical to previously defined penalties, but the penalty below *∊* corresponds to a linear continuation, which monotonically increases with decreasing determinant. In order to obtain a 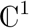-continuous penalty, the slope of this linear penalty function is equal to 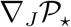 at J = ∊, and the intercept with the vertical axis at *J* = 0 is determined such that the two pieces below and above e attain the same value for *J* = ∊. Using 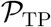avoids a separate unfolding step, and the registration task is determined by a single well-defined energy minimisation.

#### 2.5.3 Energy minimisation

The energy given by equation (6) is minimised using a conjugate gradient descent with a greedy line search. The derivatives of the penalty terms with respect to the coefficients of the SVFFD are equivalent to the derivations for the displacement-based FFD.

The gradients of the Jacobian penalties are

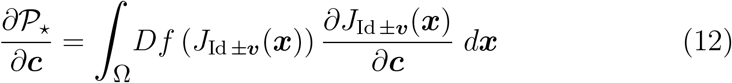

where

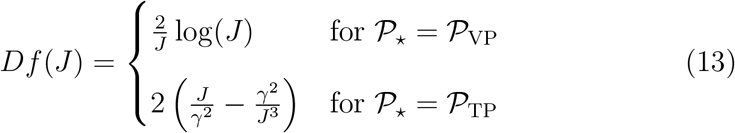

and 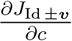is obtained using Jacobi’s formula as in Modat et al. (2012).

The derivatives of the image dissimilarity term can be written as

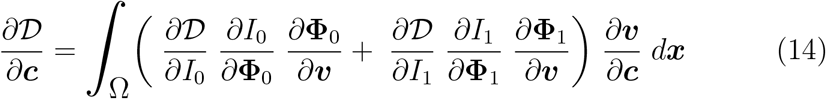

where Φ_0_ = exp(–*υ*/2), and Φ_1_ = exp(–*υ*/2). The factor in parenthesis corresponds to the sum of voxel-wise non-parametric gradients of the image dissimilarity measure with respect to a change in the SVF for each reference point *x* ∊ Ω. The partial derivatives 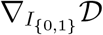 are given for NMI based on the Parzen window estimation of the joint probability distribution using cubic B-spline kernels in Modat et al. (2009), and for LNCC in Avants et al. (2008).

The numerical approximation of the derivatives of the exponential maps 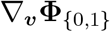 depends on the integration method. The formulas for the Euler integration are equivalent to the temporal diffeomorphic free-form deformation (TDFFD) algorithm of De Craene et al. (2012), which is a variant of LD-DMM based on a B-spline parametrisation. While more accurate than SS, a Runge-Kutta integration is too time consuming for an efficient gradient evaluation. Faster approximations based on SS are outlined in Ashburner (2007) and Modat et al. (2012). Here, we use an even simpler approximation based on the assumption that the gradient can be considered to be an instantaneous velocity field. The same is assumed by the viscous fluid method (Christensen et al., 1996) and diffeomorphic Demons (Vercauteren et al., 2009). The current SVFFD is composed with this SVF using the Baker-Campbell-Hausdorff (BCH) formula (c.f. Bossa et al., 2007; Schuh et al., 2014), i.e.,

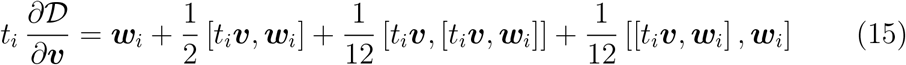

where *t*_0_ = −0.5, *t*_1_ = 0.5, and 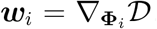. This computation is similar to log-domain Demons (Vercauteren et al., 2008). By interpolating all vector fields (including the Lie bracket [•, •]) by cubic B-spline functions defined on the same lattice as *υ*, the approximated image dissimilarity gradient with respect to *c* is given directly by the spline coefficients of the right-hand side of equation (15). Following Lorenzi et al. (2013), we use the zeroth order approximation of the BCH formula. The coefficients of *w*_i_ are obtained via convolution of 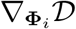with the cubic B-spline kernel (Modat et al., 2010).

### 2.6 Spatio-temporal atlas construction

Our proposed spatio-temporal atlas construction is illustrated in figure 3. In this schematic drawing, the horizontal axis corresponds to the temporal domain and the image planes orthogonal to it depict the 3D space of the atlas. Each image plane pictures an atlas template to which a given sample brain image acquired at time *t_i_* is registered to, in order to infer the transformation which relates it to the atlas space. The temporal density of the observations is therefore illustrated by the spacing of the cross-sectional image planes along the time axis. The adaptation of the kernel width to this age distribution is shown by three exemplary Gaussian curves overlaid over the diagram.

**Figure 3.**
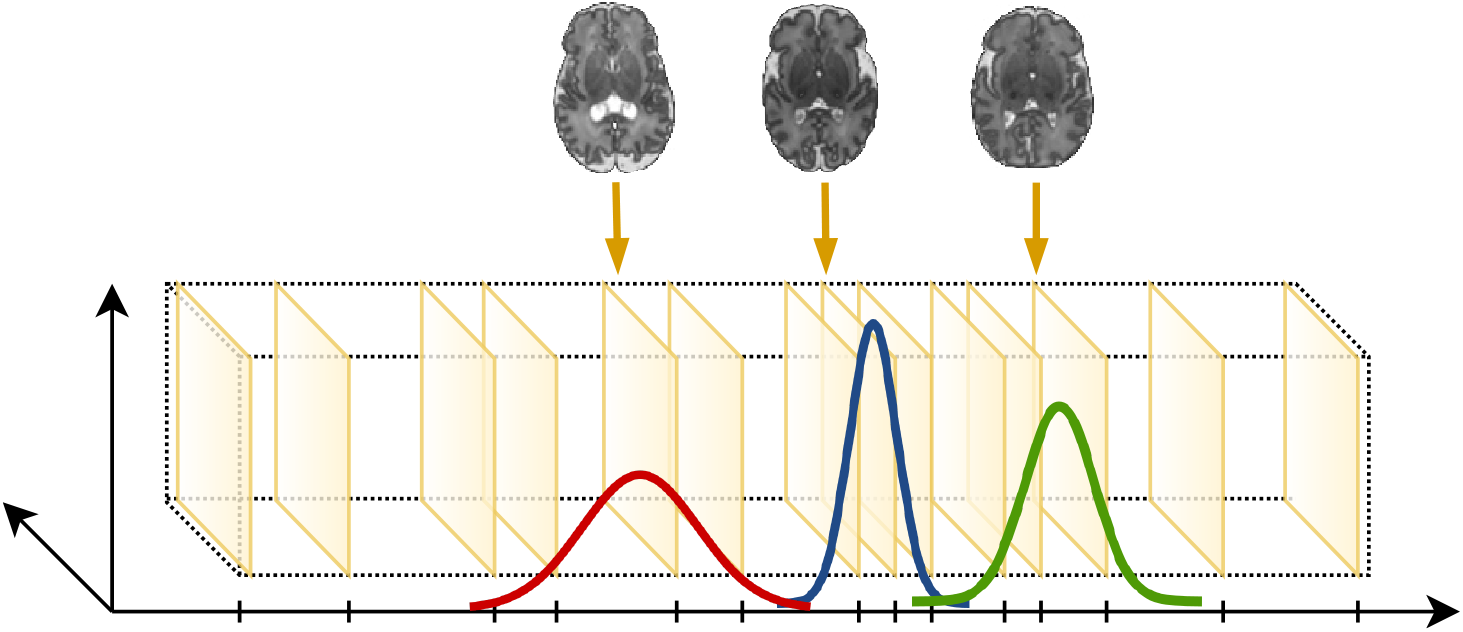
Illustration of atlas construction with time corresponding to the horizontal axis, and cross-sectional image planes depicting the spatial domain. Exemplary regression kernels are shown as overlaid curves. Arrows illustrate the subject-to-atlas mappings.

Once the recursive mean intensity function in section 2.6.1 is defined, we modify this definition in section 2.6.2 to also incorporate an estimate of the longitudinal deformations of the atlas space from one time point to another. The steps which are iterated to construct a spatio-temporal atlas from *n* neonatal brain images of different ages are then summarised.

#### 2.6.1 Recursive definition of spatio-temporal atlas

The mean intensity and brain shape template at iteration *k* is given by

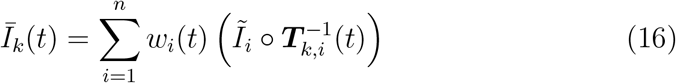

where 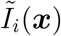is the *i*-th globally normalised image, and *T_k,i_* is the current estimate of the diffeomorphism relating it to the spatio-temporal atlas space.

The weights used for temporal regression are normalised Gaussian kernels centred at time point *t*, with age-dependent standard deviation σ*_t_*, i.e.,

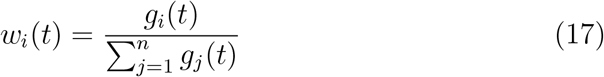

where

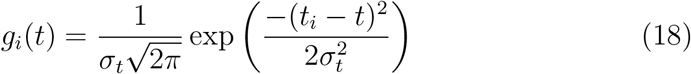

These are truncated below a small threshold to limit their global extent. An adaptive kernel width has been shown by Serag et al. (2012a) to be advantageous over a fixed kernel width for the construction of a spatio-temporal atlas of the early developing brain. This is due to the age at scan distribution of most neonatal datasets, where neonates are ideally imaged close to birth and most are born around term between 37 to 40 weeks gestation. Adapting the kernel width to be narrower at term, combined with a data selection which eliminates sharp peaks in the age distribution, effectively smooths the age distribution and makes it more uniform. This results in temporally more consistent templates, given a more even distribution of anatomical samples. The weighted average given by equation (16) shifts the barycentre of deformed brain anatomies towards those samples that are closer in age to the atlas time point. Images of an earlier or later developmental phase have less or no influence. This is an important property given the large shape differences at different ages, and the aim to construct an atlas that resembles the mean brain shape at each stage of brain development during early life.

The subject-to-atlas deformations are modelled as the composition

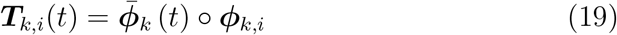

where

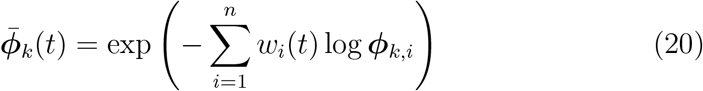

is the weighted Log-Euclidean mean (Arsigny et al., 2006) of inverse subject-to-atlas deformations obtained through SVFFD registration of *Ĩ_i_*to the age-matched average image of the previous iteration, i.e., *Ĩ*_*k*−1_(*t_i_*). The logarithmic maps are hereby given directly by the computed SVFs. The cross-sectional diffeomorphism mapping *Ĩ_i_*into the atlas space at *t_i_* is given by

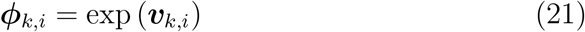

where

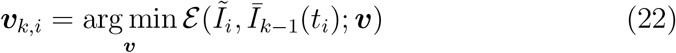

is the (local) minimum of the energy given by equation (6), and υ_0,*i*_ = 0.

#### 2.6.2 Joint estimation of mean shape and growth

In the previous section, the transformations relating each brain image to different time points were considered to have the same anatomical co-domain. This neglects the sequence of residual mean deformations 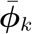 that deform the atlas space at each time point differently for the mean shape to flow towards the weighted barycentre of the nearby observations at the respective age. As illustrated in figure 4, we can track the sequence of space deformations applied at each time *t*, and use these to derive a longitudinal coordinate map, which relates an observed age *t*_i_ to any other time point in order to correct for this anatomical mismatch in the co-domain, i.e.,

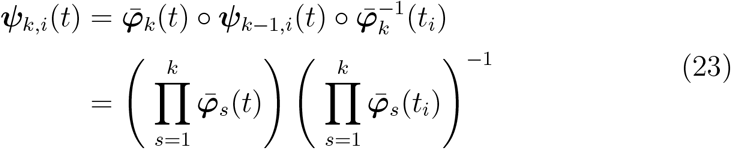

where Π denotes a sequence of function compositions, and 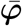are the adjusted Log-Euclidean mean transformations defined below. The non-recursive definition follows from the assumption that the initial longitudinal deformation between time *t_i_* and any other time point is the identity transformation, i.e., *ψ*_0,i_(*t*) = Id. Because each transformation is the exponential map of a SVF, all compositions can be approximated using the BCH formula, such that the composite transformation is another SVFFD.

**Figure 4.**
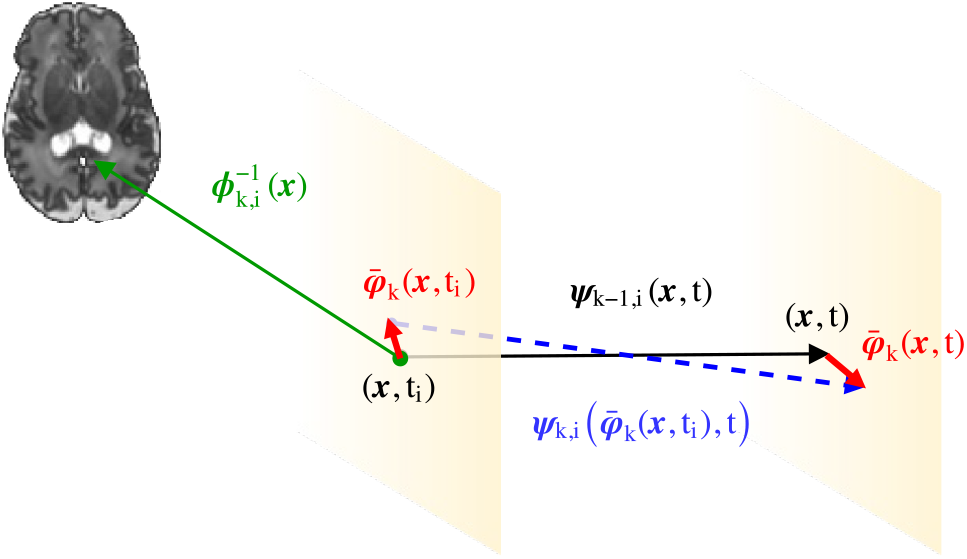
Recursive estimation of mean growth via composition of space deformations. The cross-sectional SVFFD *ϕ_k,i_* (green arrow), computed by non-rigid registration of the *i*-th image to the atlas template at age *t_i_* (left plane), is composed with the current longitudinal deformation *ψ_k−1,i_* (black arrow) to map the image to time *t* (right plane). This longitudinal estimate of mean growth evolves together with the spatial anatomical coordinates at the two time points which are deformed at iteration *k* by the age-specific mean deformations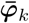 (red arrows), resulting in the new estimate *ψ_k,i_* (blue arrow).

The composition of the subject-to-atlas transformation given by (21), which is obtained by registering the *i*-th image to the template *Ī_k-1_(*t*_i_*), with the longitudinal deformation (23), yields an age-specific deformation, i.e.,

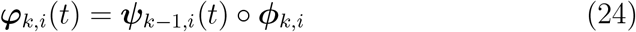

For a set of discrete time points, these transformations may be found by separate registrations of the *i*-th image to the different atlas templates as in Gholipour et al. (2017). However, obtaining it instead via composition with an estimate of the longitudinal atlas deformation, explicitly enforces temporal consistency and reduces the number of required registrations to *n*.

The residual atlas deformation is now given by the Log-Euclidean mean of the age-adjusted atlas-to-subject transformations. It is thus given by

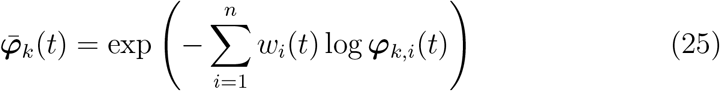

which are further used to redefine the total age-dependent deformations, i.e.,

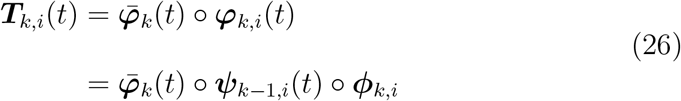

This concludes the definition of the coordinate maps involved in this new recursive spatio-temporal atlas construction based on the diffeomorphic SVFFD algorithm and the Log-Euclidean mean. As mentioned, all function compositions are efficiently approximated using the BCH formula, resulting in a compact representation of all coordinate maps. Moreover, while the mean intensity and shape template can be evaluated for any time *t*, during the atlas construction, only those images corresponding to the discrete time points *t*_i_ are needed. The result of the atlas construction is not only the spatio-temporal atlas of mean shape and intensity, and the transformations mapping each anatomy into the atlas space, but also an estimate of the longitudinal deformations corresponding to mean growth. It should be noted, though, that these deformations primarily reflect coarse anatomical changes, which is in part due to the approximations and the remaining misalignment after deformable registration, but also because finer details are the result of the temporal kernel regression of intensities at each atlas coordinate. The final mean shape is a combination of both, the sequence of Log-Euclidean mean deformations, and the regression of the intensities.

In summary, the atlas construction iterates the following steps:

1. Generate |{*t_i_*}| ≤ *n* template images *Ī*_k-1_(*t_i_*) given *Ĩ_i_* and *T_k−1,i_* (16).
2. Compute *ϕ_k,i_* = exp(υ_*k,i*_) from *i*-th image to template at *t_i_* (21).
3. Compose maps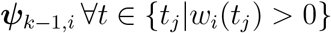 given *ψ*_*k*−2,*i*_ and 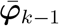(23).
4. Compose maps *ϕ_k,i_* with 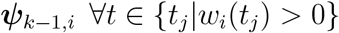(24).
5. Compute Log-Euclidean means 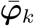 for each observed time *t*_i_ (26).

After about 5-10 iterations (or convergence based on residual deformations), template images of mean intensity and shape can be generated for a set of discrete time points together with the longitudinal deformations *ψ* between consecutive templates. The mean growth deformations can be used to initialise a subsequent longitudinal registration of the atlas time series, for example, using the TDFFD method (De Craene et al., 2012) with initial velocities given by the SVFs of the SVFFDs.

## 3. Results

### 3.1 Evaluation criteria

A number of measures were used to assess the quality of the pairwise registrations and the alignment of the brain anatomies in the atlas space. A more accurate alignment directly translates into sharper brain templates. Different subsets of criteria are used to select registration parameters, and to compare atlases built with different approaches. First, we define the per-voxel/-structure measures, before detailing the calculation of the summarising average measures.

#### 3.1.1 Label entropy

A segmentation-based voxel-wise measure for evaluation of spatial normalisation procedures was proposed by Robbins et al. (2004). For this, the entropy of the labels assigned to a point in the standard space is evaluated. When all labels are in agreement, i.e., the atlas label is uniquely determined, the entropy is zero. It attains its maximum when all labels are equally likely, i.e., when each segmentation suggests a different label for a given atlas coordinate. The label entropy at atlas point *x* at time point *t* is given by

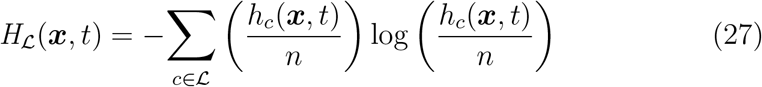

where *h_c_ (*x,t**) is the number of times class label *c* is propagated from one of the brain segmentations by their respective image-to-atlas transformations using nearest neighbour interpolation, and 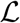 denotes the set of integral labels, i.e., either the set of Draw-EM tissue classes 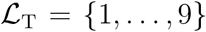 or the set of Draw-EM brain structure labels 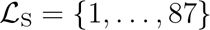. We refer to the entropy of the tissue class labels as tissue (class) entropy, and the entropy of the whole brain structure labels as structure (label) entropy, respectively.

#### 3.1.2 Intensity entropy

The entropy of intensity samples at atlas coordinate (*x,t*) quantifies the uncertainty of the mean intensity estimate at this point, i.e.,

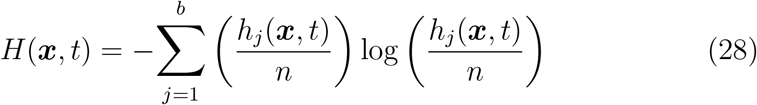

where a histogram 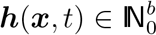 with *b* bins is used to estimate the probability distribution of the normalised intensities. The more diverse the intensities are, the more uncertain is the mean estimate.

#### 3.1.3 Standard deviation

Another measure of voxel-wise intensity variation, also used by Gholipour et al. (2017) to quantify atlas sharpness, is the standard deviation, i.e.,

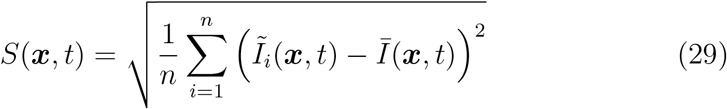

The lower this value, the lower is the standard error of the mean intensity.

#### 3.1.4 radient magnitude

The magnitude of the gradient of mean intensities in the atlas space, i.e.,

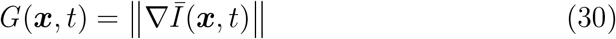

where ∇*I* is approximated using central differences, is proportional to the sharpness of image edges. When boundaries of structures with distinct appearance are aligned well, we observe a strong gradient at this boundary.

This criterion was also used by Gholipour et al. (2017). To specifically quantify the sharpness of the cortex in the mean images, we assess the gradient magnitude of the cGM probability map of the atlas.

#### 3.1.5 Pairwise overlap

The overlap measure quantifies the accuracy of the pairwise registrations using the automatic whole brain segmentation. Given the number of true positive (TP), true negative (TN), false positive (FP), and false negative (FN) label assignments for each structure and pair of segmentations of the *i*-th and *j*-th images propagated to the atlas at time point *t_l_*, the mean Dice similarity coefficient (DSC) (Dice, 1945) of the c-th structure is given by

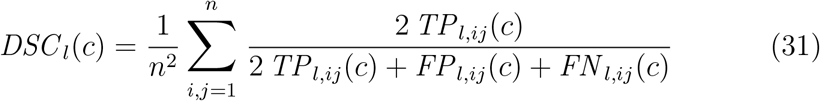

#### 3.1.6 Temporal consistency

To quantify the effect of the choice of temporal kernel width, we define a measure of temporal consistency (TC). The TC of structure *c* sampled at *n_t_* ordered time points *t_l_* (i.e., *t_l_* < *t*_*l*+1_) is the generalised DSC given by

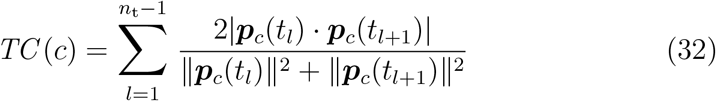

where the scalar product of two probabilistic segmentations is given by

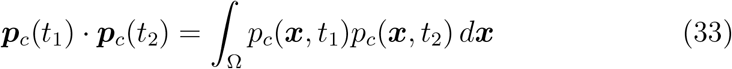

and the squared *ℓ*_2_-norm is 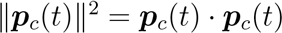.

#### 3.1.7 Average of spatio-temporal measures

The voxel-wise measures are averaged for each discrete atlas time point *t_l_* using the probabilistic segmentation of the atlas into nine Draw-EM classes. Given a quality measure *Q(**x***, *t*), the average for 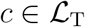 and *t* = *t_l_* is

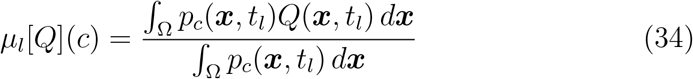

### 3.2 Selection of model parameters

#### 3.2.1 Image dissimilarity measure

For the construction of a neonatal brain atlas using our previous pairwise construction techniques (Serag et al., 2012a; Schuh et al., 2014) and the new groupwise approach presented in section 2.6, we first compared registration results between all pairs of images for three subsets of ten brain images each (representing different phases of brain development) for two different commonly used dissimilarity measures, namely NMI and LNCC. In the work of Gholipour et al. (2014) on constructing a fetal brain atlas using SyGN (Avants et al., 2010), the use of LNCC resulted in slightly sharper templates. In our case, for both the FFD method (Rueckert et al., 1999) used in the original approach of Serag et al. (2012a) and our proposed SVFFD algorithm, mean overlap of brain structures was found to be consistently higher for NMI when compared to the results obtained with LNCC. A registration using NMI is also more efficient (on average less than 40% of CPU time). Given the better performance and lower computational cost of NMI, we used it for all pairwise neonatal brain image registrations. Note that NMI has also been used for the construction of previous spatio-temporal neonatal atlases (Kuklisova-Murgasova et al., 2011; Serag et al., 2012a; Schuh et al., 2014).

#### 3.2.2 Control point spacing and regularisation weights

With the image dissimilarity measure fixed, we performed an exhaustive grid search over the regularisation weights and different control point spacings using the subset of images used to select the dissimilarity measure. For the reference method of Serag et al. (2012a) based on the displacement-based FFD model, a control point spacing of 2.5 mm resulted in the best compromise between overlap and amount of folding. In case of the diffeomorphic SVFFD model, a smaller control point spacing of 2 mm was selected.

A comparison of the evaluation measures for both models with the selected parameters, and evaluated for the three subsets of in total 30 neonatal subjects scanned at different PMAs, is given in figure 5. This first comparison of separate atlases constructed without temporal kernel regression for three time points using our previously proposed pairwise construction techniques (Serag et al., 2012a; Schuh et al., 2014) suggests that both registration methods achieve similar segmentation overlap using the selected parameters, but that the diffeomorphic SVFFD registration combined with the Log-Euclidean mean produces better atlas templates, i.e., suggesting an advantage of our inverse-consistent and diffeomorphic atlas construction presented in Schuh et al. (2014) over the original approach proposed by Serag et al. (2012a).

**Figure 5.**
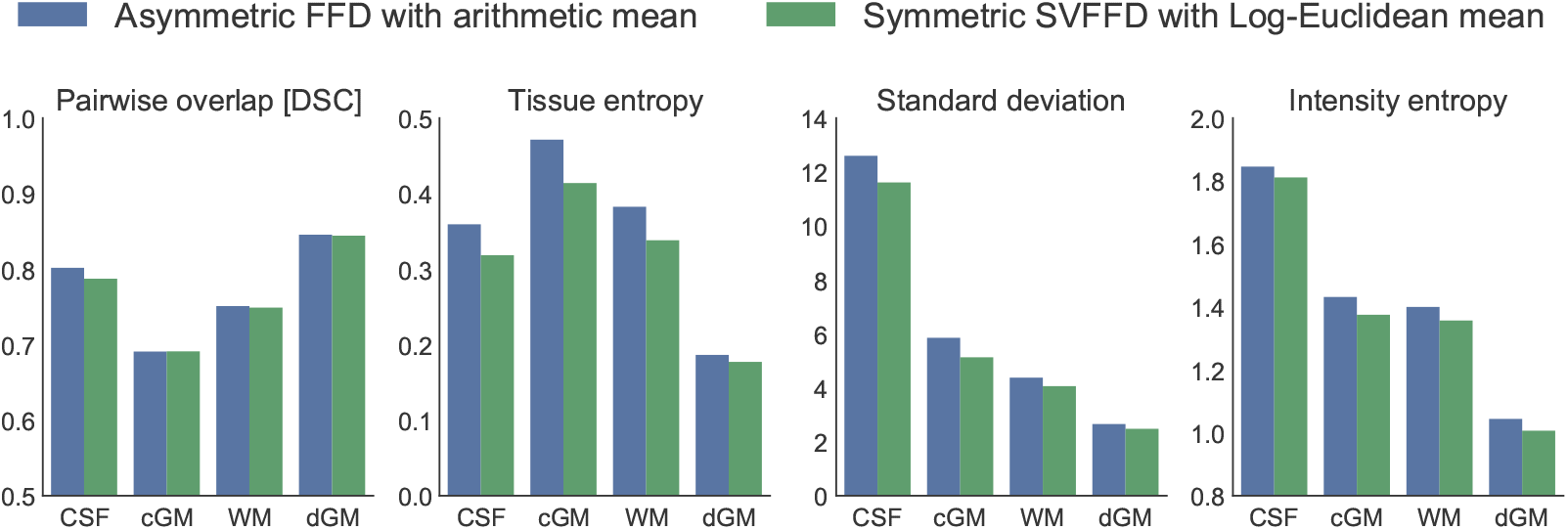
Comparison of evaluation measures for different pairwise atlas constructions based on a subset of in total 30 neonatal subjects used for parameter selection. This comparison suggests that, although overlap between image pairs is similar, the template created with the SVFFD model and Log-Euclidean mean is slightly sharper.

Having determined the registration parameters for constructing a spatiotemporal neonatal atlas using direct registrations between all pairs of images, we performed a second parameter search for our new groupwise approach based on previously determined parameter ranges. This was done, because at each iteration the target image is the current average brain image, which has a different appearance than an actual brain scan. For this groupwise parameter search, we constructed a single time point atlas at 40 weeks PMA from a subset of 64 brain images of term-born neonates. The SVFFD parameters chosen for the pairwise atlas construction were found to be suitable also for the groupwise method. Due to the iterative refinement, the approach demonstrated to be more robust towards variations in parameters. However, the initial target images after affine alignment contain noticeable artefacts due to a significant amount of residual anatomical misalignment. We therefore constrain the deformations at early iterations more strongly using a sparser control point placement, and progressively half the control point spacing of the SVFFD after a few iterations. For the first two iterations, a control point spacing of 8 mm at the highest image resolution level is used. At iterations three to five, the control point spacing is 4 mm, whereas it is 2 mm for the final iterations. We found that the quality of the atlas did not substantially improve after 8 iterations, as can be seen in figure 6.

**Figure 6.**
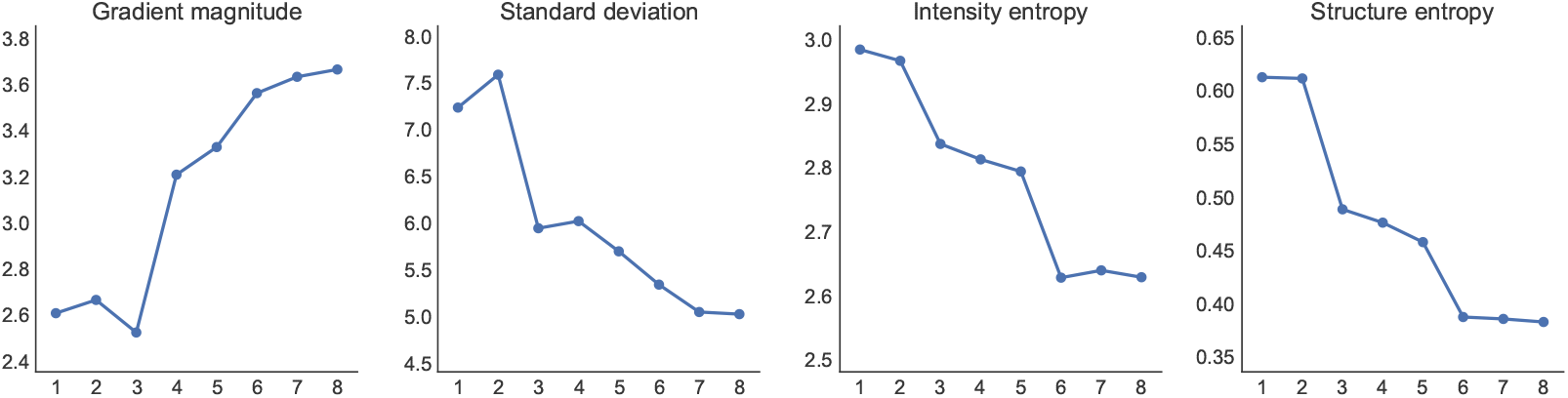
Convergence of atlas quality measures in case of groupwise construction. The horizontal axis corresponds to number of completed iterations, *k*. The jumps from iteration *k* = 2 to *k* = 3, and from *k* = 5 to *k* = 6 are the result of halving the minimum control point spacing of the SVFFD at the highest image resolution level between iterations.

#### 3.2.3 Standard deviation of temporal regression kernel

For the construction of mean images at weekly intervals of the age range from 36 to 44 weeks PMA, for which a sufficient number of brain scans is available in our dataset, we use an algorithm similar to Serag et al. (2012a) to determine an age-specific standard deviation, *σ_t_,* of the temporal regression kernel for each age *t_l_* ∈ {36,37,38,39,40,41,42,43,44}. Our kernel width selection differs from the algorithm proposed by Serag et al. (2012a) in that no stopping criterion based on the difference in brain volumes is considered, because our goal is a uniform distribution of brain samples at the equally spaced discrete time points. We first specify a constant kernel width, *σ,* and determine the median number, *m*, of images with non-zero kernel weight for each discrete atlas age *t_l_*. We then iteratively increase or decrease each kernel width *σ_tl_* by a small value until the number of images is equal to *m*.

Using this algorithm, we determined adaptive kernel widths for different target *σ* values, and constructed a separate spatio-temporal atlas for each corresponding set of kernel widths listed in table 2, and visualised in figure 7, using our previous approach presented in Schuh et al. (2014). The TC of tissue classes for each constructed atlas is shown in figure 8. As expected, it is observed that TC improves with increasing kernel widths. When *σ →* ∞, the atlas becomes constant in time, and thus has maximum TC, but fails to model temporal shape differences. Reducing *σ* results in more distinct and sharper mean brain shapes for each time point, with fewer brain images contributing to each template image. We therefore also compared the average pairwise overlap of spatially normalised Draw-EM structures and the mean voxel-wise standard deviation of T2w image intensity samples in figure 9. We expect an atlas with a consistent quality across time to achieve a similar DSC value for each time point, without sacrificing a substantial reduction of overall quality. We should, however, also consider the higher anatomical detail at later time points due to increased cortical folding.

**Table 2.** Adaptive kernel widths.

| $\sigma$ | 0.25 | 0.50 | 0.75 | 1.00 |
| --- | --- | --- | --- | --- |
| $m$ | 44 | 101 | 143 | 196 |
| $\sigma_{36}$ | 0.51 | 0.99 | 1.31 | 1.64 |
| $\sigma_{37}$ | 0.35 | 0.69 | 0.99 | 1.31 |
| $\sigma_{38}$ | 0.25 | 0.50 | 0.75 | 1.00 |
| $\sigma_{39}$ | 0.14 | 0.39 | 0.56 | 0.77 |
| $\sigma_{40}$ | 0.14 | 0.31 | 0.43 | 0.64 |
| $\sigma_{41}$ | 0.11 | 0.27 | 0.43 | 0.73 |
| $\sigma_{42}$ | 0.11 | 0.35 | 0.56 | 0.86 |
| $\sigma_{43}$ | 0.27 | 0.51 | 0.77 | 1.18 |
| $\sigma_{44}$ | 0.51 | 0.82 | 1.09 | 1.50 |

**Figure 7.**
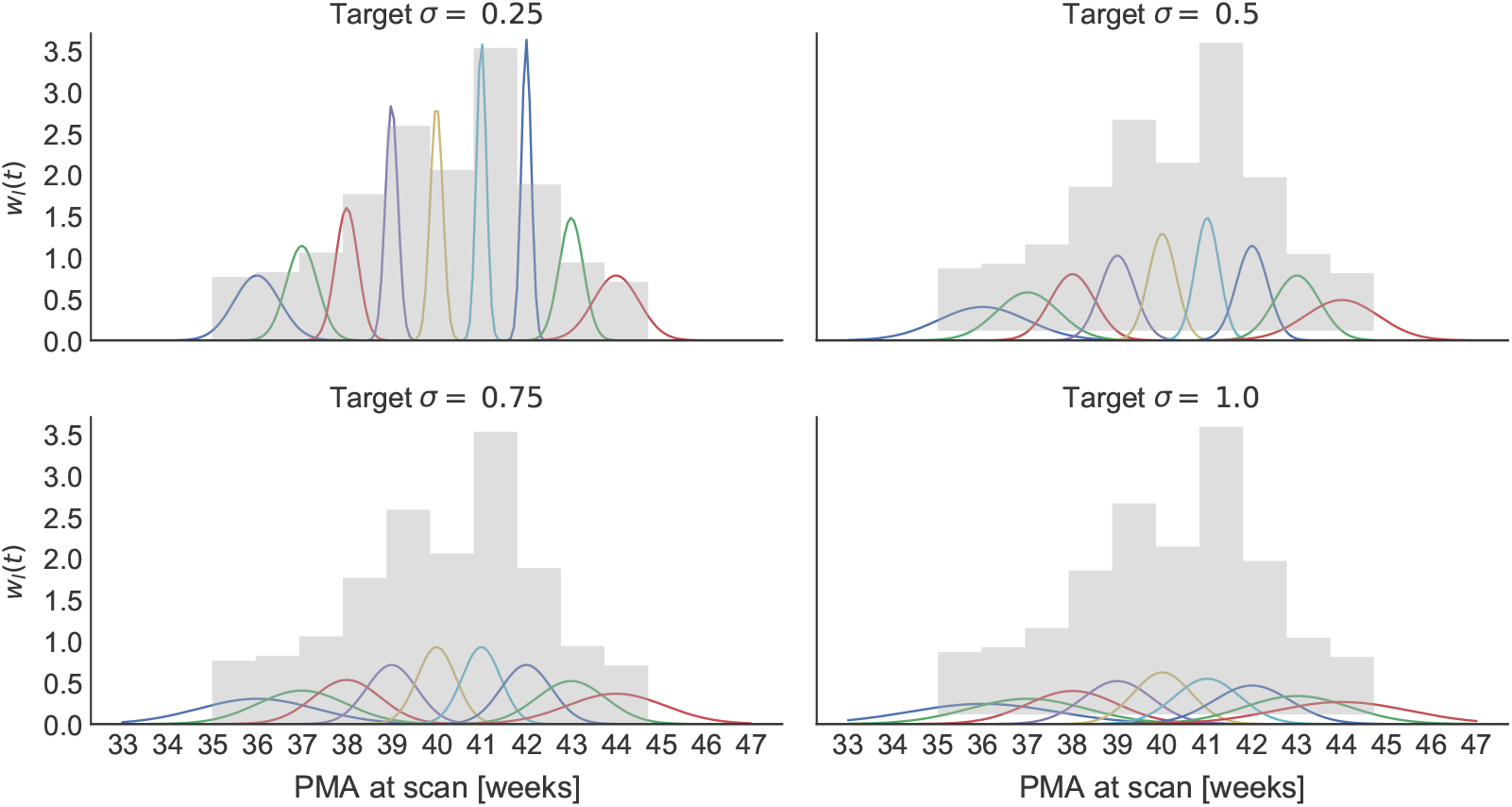
Temporal regression kernels for different ages and target σ values. The respective standard deviations of the shown regression kernels centred at the discrete time points *t_l_* are listed in table 2. The curves are overlayed on the PMA at scan histogram.

**Figure 8.**
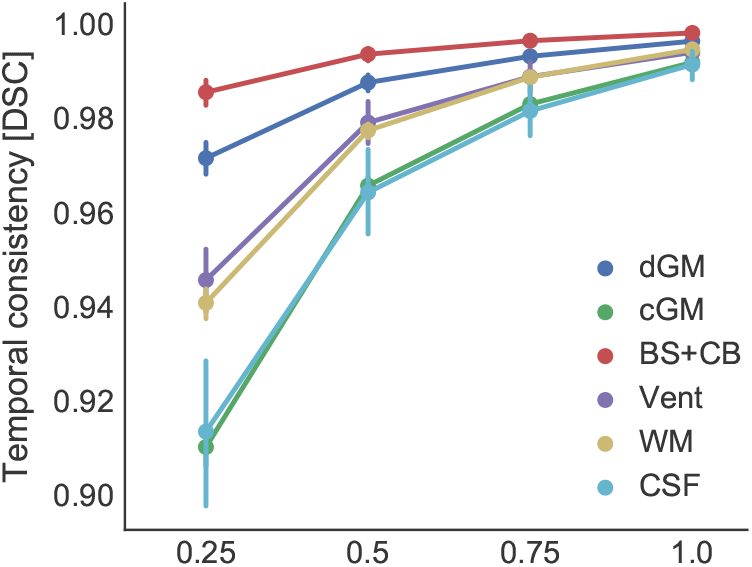
TC for different target σ values.

**Figure 9.**
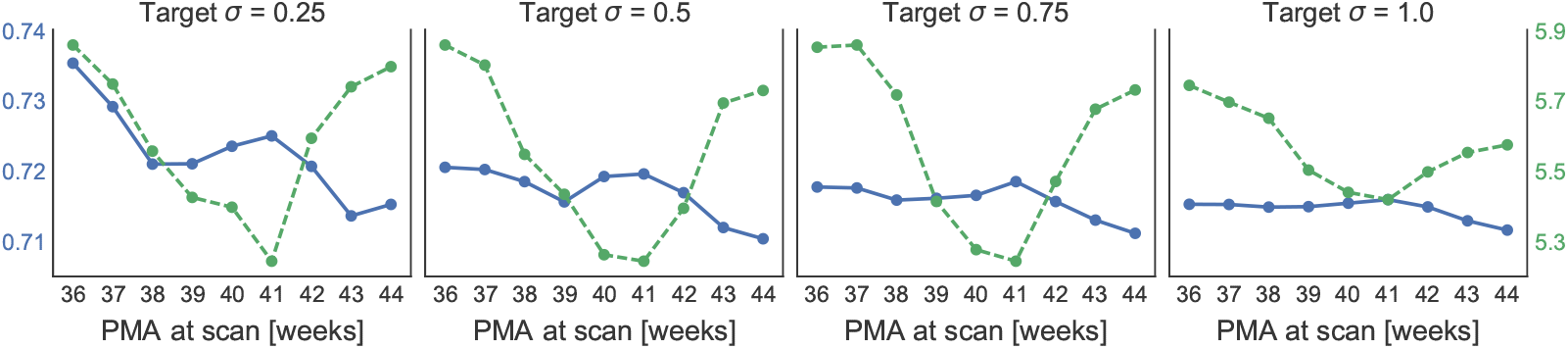
Quality measures for discrete time points of atlas constructed with approach proposed in Schuh et al. (2014) and different target σ values. The left vertical axis corresponds to the blue curve of the pairwise overlap (DSC), whereas the right axis corresponds to the dotted green curve depicting the mean voxel-wise standard deviation of T2w intensities.

A target *σ* value below half a week results in insufficient overlap to ensure temporal smoothness of the mean brain images at weekly intervals. A target kernel width of 0.75 to 1 week produces temporally smooth atlases with noticeable temporal shape differences. After visual inspection, we found that a good compromise for our dataset is obtained for target *σ* = 1. The kernel width *σ_t_* for any time point is obtained by linear interpolation, i.e.,

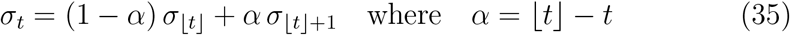

### 3.3 Comparison of atlas construction methods

With the selected parameters, we constructed a spatio-temporal neonatal atlas from 36 to 44 weeks PMA using the discussed construction methods, and evaluated these at weekly intervals. We compared the atlases constructed with our proposed approaches based on the SVFFD algorithm and Log-Euclidean mean deformations to the original method of Serag et al. (2012a), which uses the classic FFD algorithm and arithmetic mean of displacements. Our previous method proposed in Schuh et al. (2014) is closely related to the approach of Serag et al. (2012a). Both derive the transformations relating each observation to the atlas space from transformations between each unique pair of brain images. This construction technique is thus referred to as “pairwise” method, while the new groupwise construction presented here is referred to as “groupwise” approach. A key difference between our “pairwise” approach and the original “reference” method of Serag et al. (2012a) is, that we utilise topology preserving deformation models, and the Log-Euclidean mean instead of the arithmetic mean of the pairwise mappings.

To reduce the residual anatomical misalignment in the atlas space further, we additionally performed a separate iterative refinement of the template brain images at discrete time points constructed with the “pairwise” approach. For this, the mean diffeomorphisms relating each subject to the atlas space at a specific time point are refined by registering each brain image to the respective mean intensity image. The new subject-to-atlas mapping is then given by the composition of this new transformation with the Log-Euclidean mean, i.e., similar to equation (19). Given these refined subject-to-atlas transformations, a new template image is computed, and the process is repeated. This iterative approach is comparable to the spatio-temporal fetal atlas construction of Gholipour et al. (2017), but using our SVFFD algorithm and the Log-Euclidean mean instead of SyGN. We compare this “refined” atlas to our proposed “groupwise” approach. A summary of the compared construction methods is given in table 3.

**Table 3.**
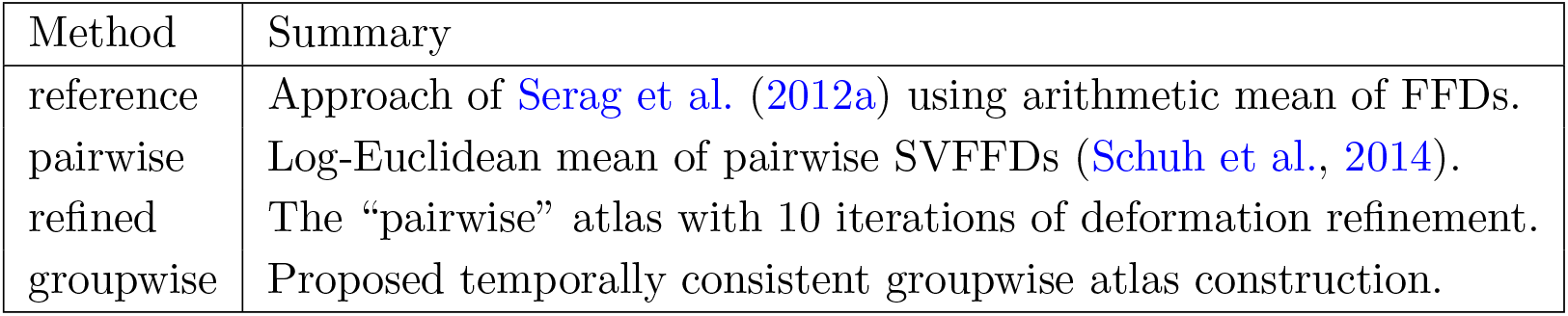
Summary of compared spatio-temporal atlas construction methods.

The mean of the voxel-wise measures averaged separately for each time point for each method over the entire brain masks is reported in table 4. Box plots of these measures visualising also the temporal variance of the average measures are shown in figure 10a, whereas plots of individual average measures over time are shown in figure 10b. The latter visualisation highlights the temporal smoothness even for average measures with a larger interquartile range, and explains this larger variance of average measures by the difference between early and late atlas time points.

**Table 4.** Mean quality measures of constructed spatio-temporal atlases. For each evaluation criterion, the result for the atlas which is most favourable is highlighted in bold.

|  | reference | pairwise | refined | groupwise |
| --- | --- | --- | --- | --- |
| Gradient magnitude | 2.837 | 3.096 | 3.648 | **3.661** |
| Standard deviation | 6.591 | 5.563 | **4.821** | 5.017 |
| Intensity entropy | 2.933 | 2.811 | 2.693 | **2.628** |
| Tissue entropy | 0.459 | 0.384 | 0.311 | **0.297** |
| Structure entropy | 0.526 | 0.454 | 0.389 | **0.382** |
| cGM gradient magnitude | 0.074 | 0.081 | 0.102 | **0.108** |

**Figure 10.**
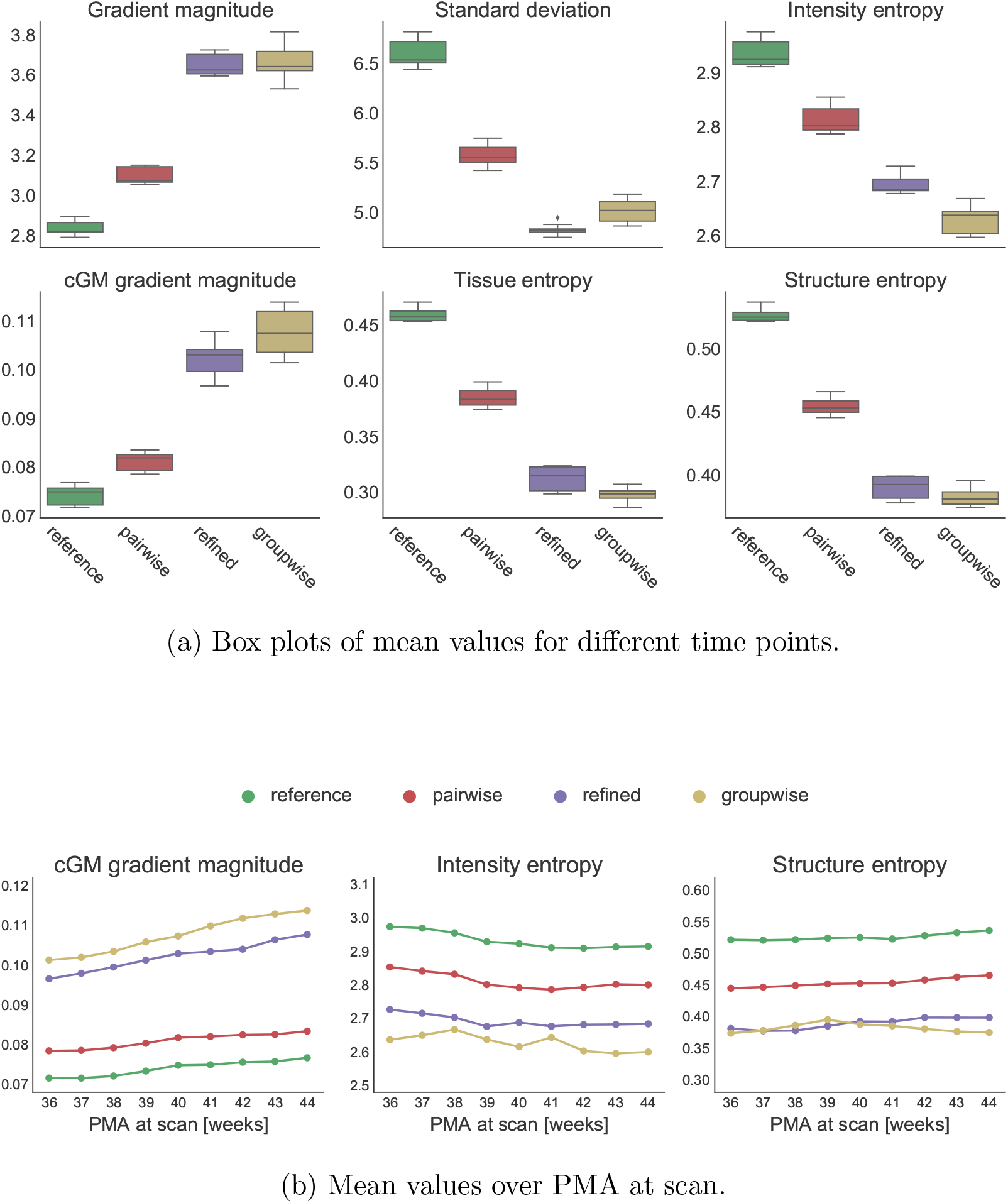
Mean quality measures of constructed spatio-temporal atlases.

As expected, a number of the evaluation measures are strongly correlated. This is especially the case for the voxel-wise label entropy of tissue classes and individual brain structures, because both are based on the automatic Draw-EM segmentations. We therefore focus on the structure entropy as it is based on a more fine granular parcellation. In regards to the intensity based evaluation criteria, the measures of intensity variance, namely standard deviation and entropy, demonstrate a similar relative performance of the construction methods. A comparison of these voxel-wise measures averaged separately for each tissue class in figure 11 shows that the apparent worse performance of the “groupwise” method when compared to the iteratively “refined” pairwise atlas is caused by a higher standard deviation of intensities for cerobrospinal fluid (CSF). This is explained by the fact that this tissue class is at the boundary of the brain mask at which the NMI similarity criterion may not be evaluated during the registration as a result of only partial overlap of the foreground regions. An improved handling of partial boundary overlap during the registration would likely improve the within CSF measures. Of higher interest is however the atlas quality for brain tissues. The average measures for cGM, WM, and deep grey matter (dGM) for each constructed spatio-temporal atlas are compared in figure 12. It can be observed that all methods perform similarly well for deep brain structures, as these require less local deformation to be aligned, but that our proposed methods improve the quality of the atlas noticeably within WM and especially cGM regions. Our pairwise atlas construction using the Log-Euclidean mean of inverse-consistent diffeomorphic mappings is more consistent near the cortex than the approach proposed by Serag et al. (2012a). This is in part because we are able to more locally align cortical features with a lower control point spacing of the SVFFD, which does not introduce folding. In contrast, the FFD model requires stronger regularisation in order to preserve the topology of the anatomies. Moreover, the iterative approaches clearly outperform the pairwise methods due to the iterative compensation of residual anatomical misalignment. Our proposed “groupwise” spatio-temporal atlas construction performs slightly better than the individual iterative refinement of atlas templates at discrete time points.

**Figure 11.**
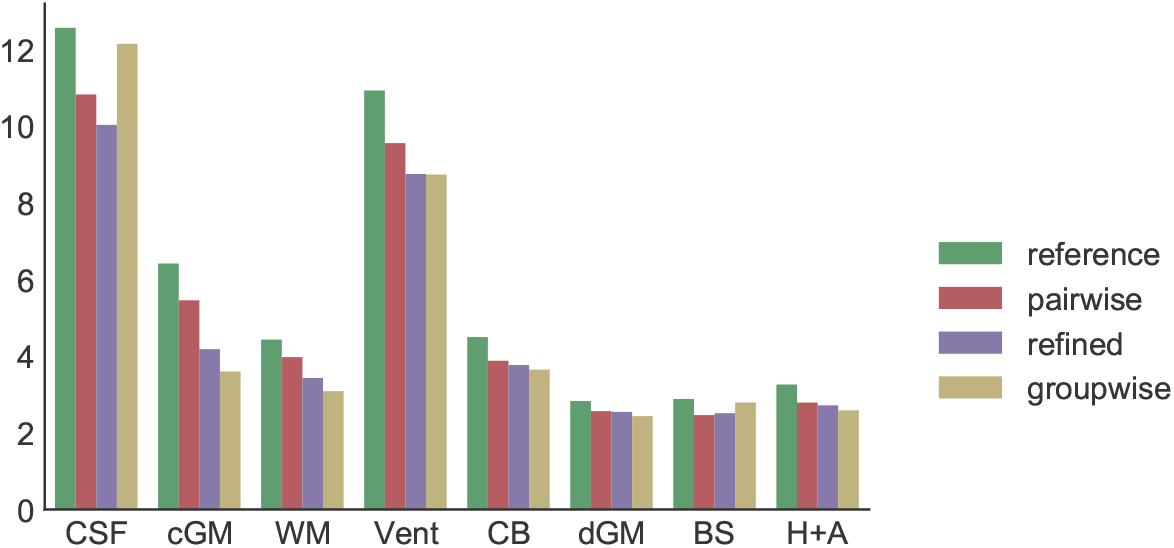
Intensity standard deviation averaged separately for CSF, cGM, WM, ventricles (Vent), cerebellum (CB), dGM, brainstem (BS), hippocampi and amygdalae (H+A).

**Figure 12.**
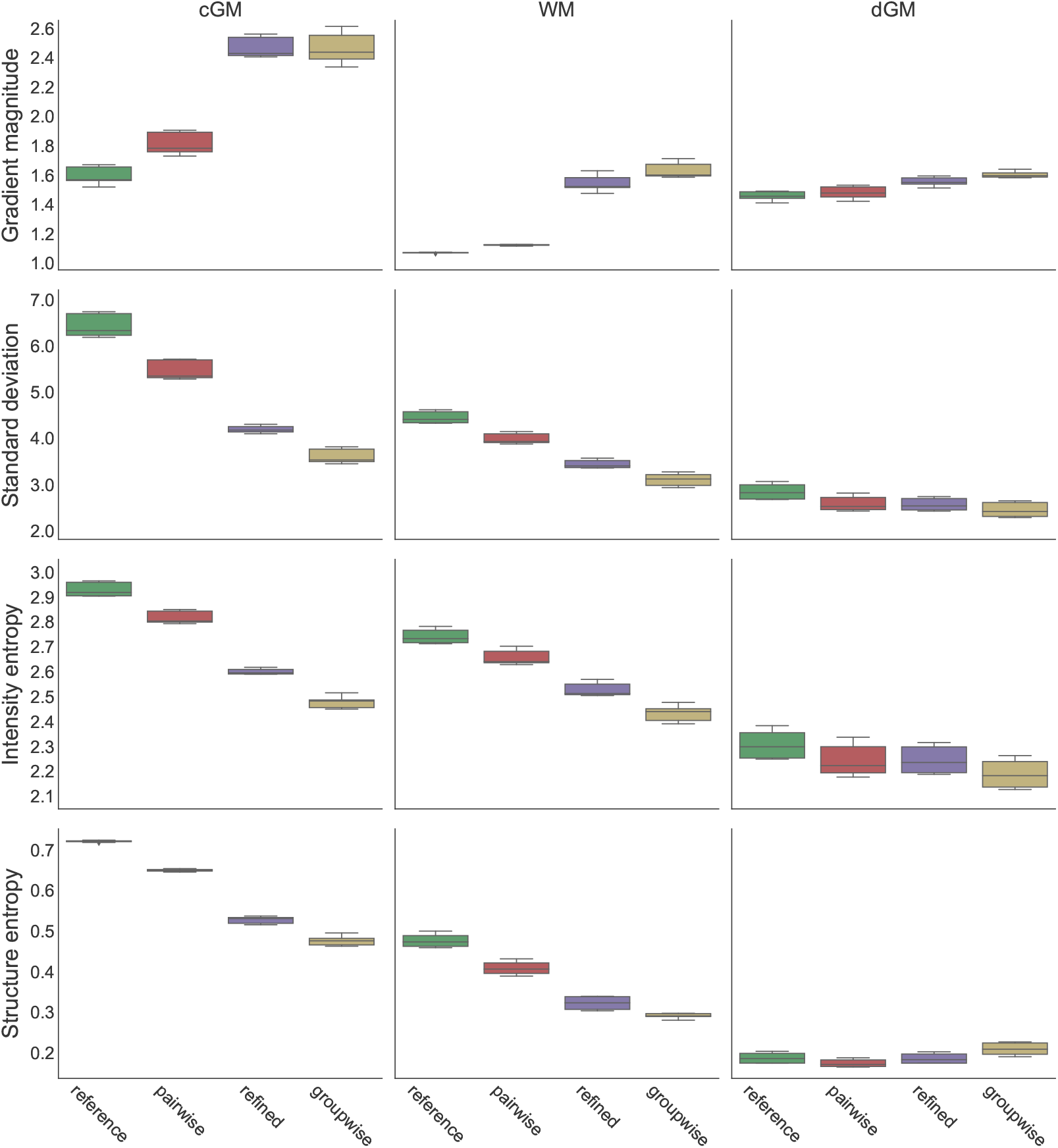
Atlas quality measures averaged separately for cGM, WM, and dGM.

### 3.4. Visual assessment of neonatal atlases

Having observed improvements in the defined quantitative measures, we further assess the quality of the constructed atlases visually. Templates for 40 weeks PMA constructed with the reference method of Serag et al. (2012a), our derived pairwise approach using the SVFFD model and Log-Euclidean mean, the template of the pairwise method after 10 iterations of individual subject-to-atlas deformation refinement, and the template generated by the proposed groupwise approach after 8 iterations are compared in figure 13a. Most noticeable are the sharper cortical details achieved by the iterative approaches. Differences in the atlases obtained with either of the two pairwise or iterative techniques, respectively, are more subtle, and less clear by inspecting a single cross-section. Slightly clearer cortical boundaries may be observed in the atlas created with our pairwise method when compared to the reference approach, in particular at the frontal lobe. These differences are more apparent when examining coloured renders of the voxel-wise intensity and tissue class entropy measures in figure 13b. Our pairwise method achieves more anatomical consistency in the alignment of the individual brain scans in the atlas space, but both methods produce atlases of high quality. An advantage of our approach is a consistent modelling of mean shape by topology-preserving deformations.

**Figure 13.**
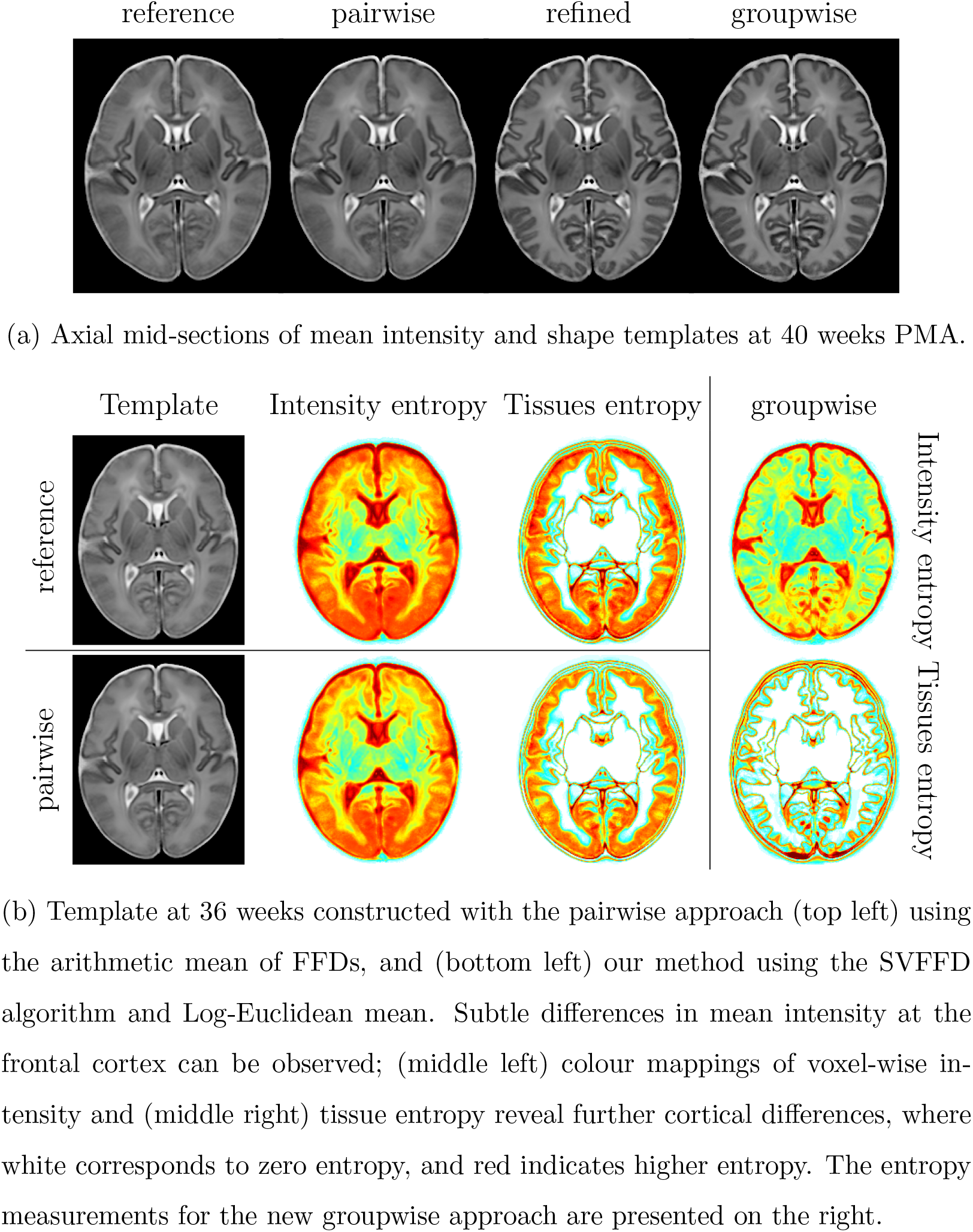
Visual comparison of templates constructed with different approaches.

The improved temporal consistency of the atlas constructed with our groupwise approach, as opposed to an individual iterative refinement of each time point, is highlighted in the detail views of smaller ROIs in figure 14. When brain scans are registered individually to each template, developing cortical structures may be missing at certain time points or some folding patterns recede instead of deepen further, which is not expected for normal development. This is not observed in the temporal sequences generated by our groupwise method, which consistently maps each brain scan to each individual time point considering the longitudinal inter-atlas deformations.

**Figure 14.**
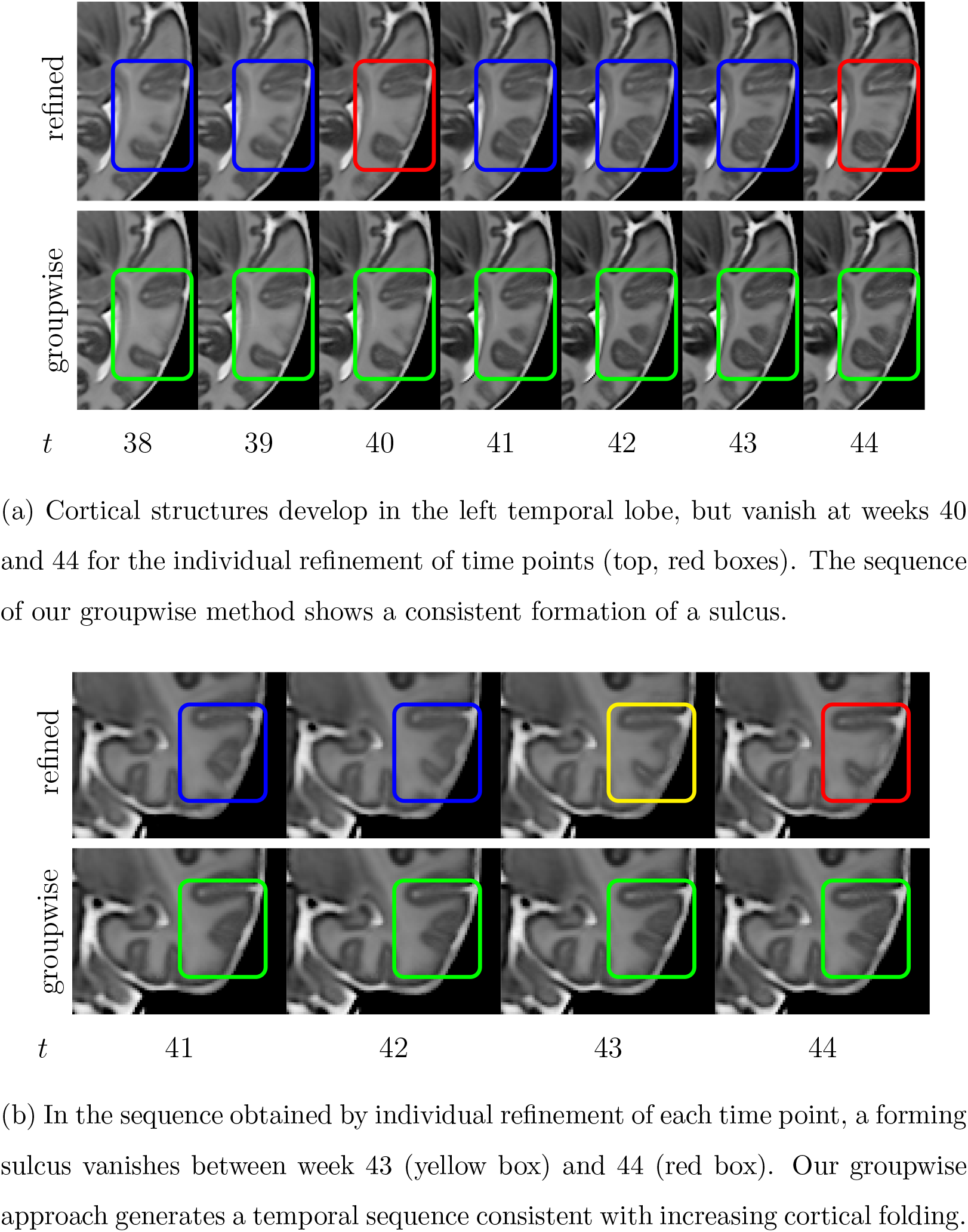
Detail views of axial cross-sections of atlases constructed (top) with the pairwise approach followed by iterative refinement, and (bottom) our groupwise method.

T1w and T2w mean shape and intensity images at weekly intervals for the age range from 36 to 44 weeks PMA generated with the proposed groupwise technique are displayed in figure 15, along with the spatial tissue probability map for cGM. The clear markedness of cortical folds is reflected in the sharpness of the spatial cGM probability distribution.

**Figure 15.**
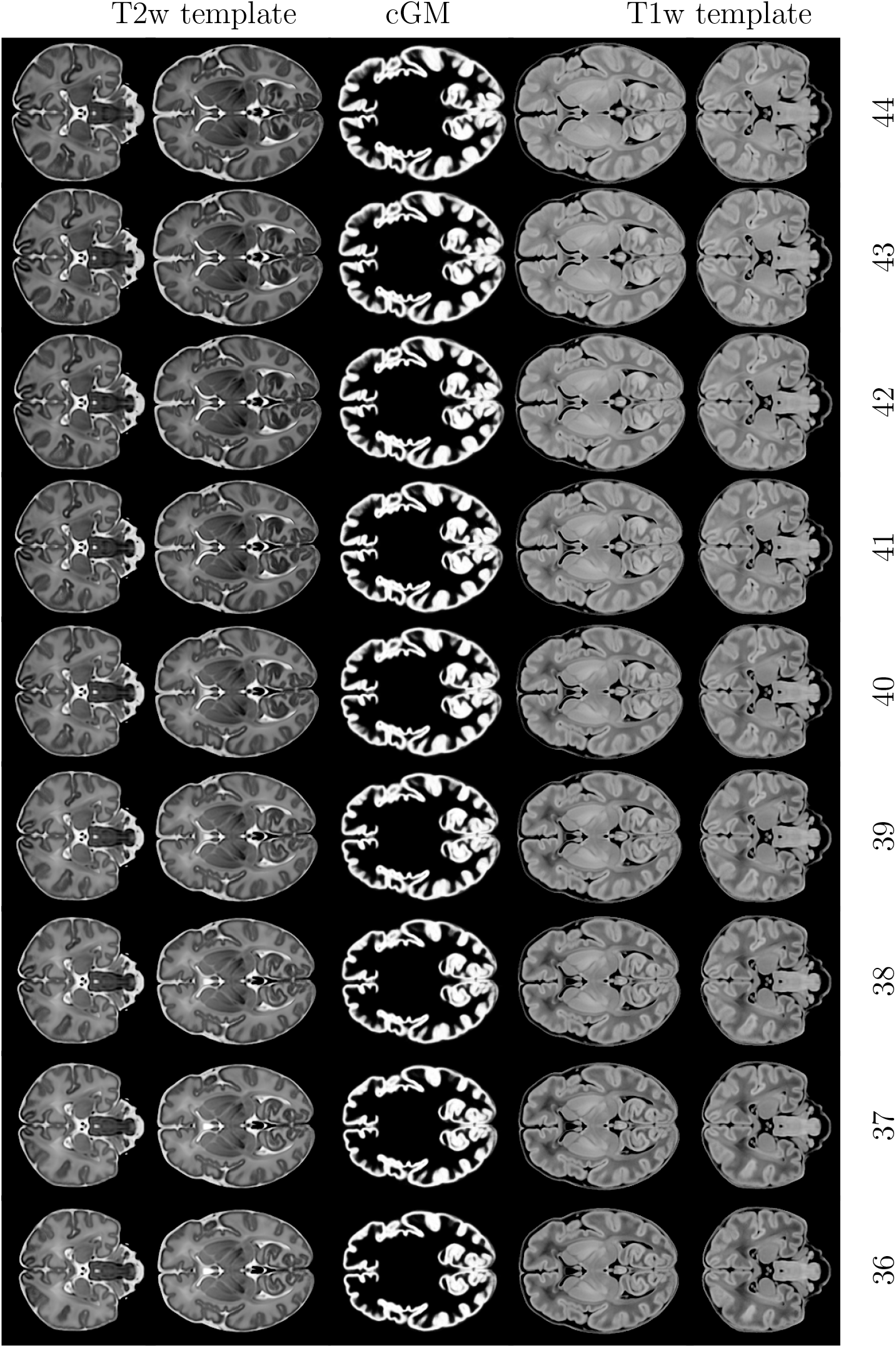
Spatio-temporal atlas constructed with the proposed groupwise approach.

### 3.5. Qualitative comparison to previous atlases

Thus far, we compared atlases constructed with our methods based on the SVFFD algorithm to the atlas built according to the approach of Serag et al. (2012a) from the same set of images, and an improved registration using the Medical Image Registration ToolKit (MIRTK). In figure 16, we give a comparison to the spatio-temporal neonatal brain atlases pre-dating our work, that were made publicly available by the original authors of these methods. The significant improvement in image quality of the new atlases is clearly noticeable. One reason for this is that the brain images acquired by dHCP are of remarkably higher quality than previous datasets of preterm-born neonates that were available for the construction of these previous atlases. Another reason is that at each time point of our new atlases, 196 anatomies are being averaged, whereas the mean images created by Serag et al. (2012a) are the average of 15-19 individual brain scans. This explains the lower cortical detail of our pairwise atlas. High quality and marked cortical folds are present in the atlas generated by our groupwise method.

**Figure 16.**
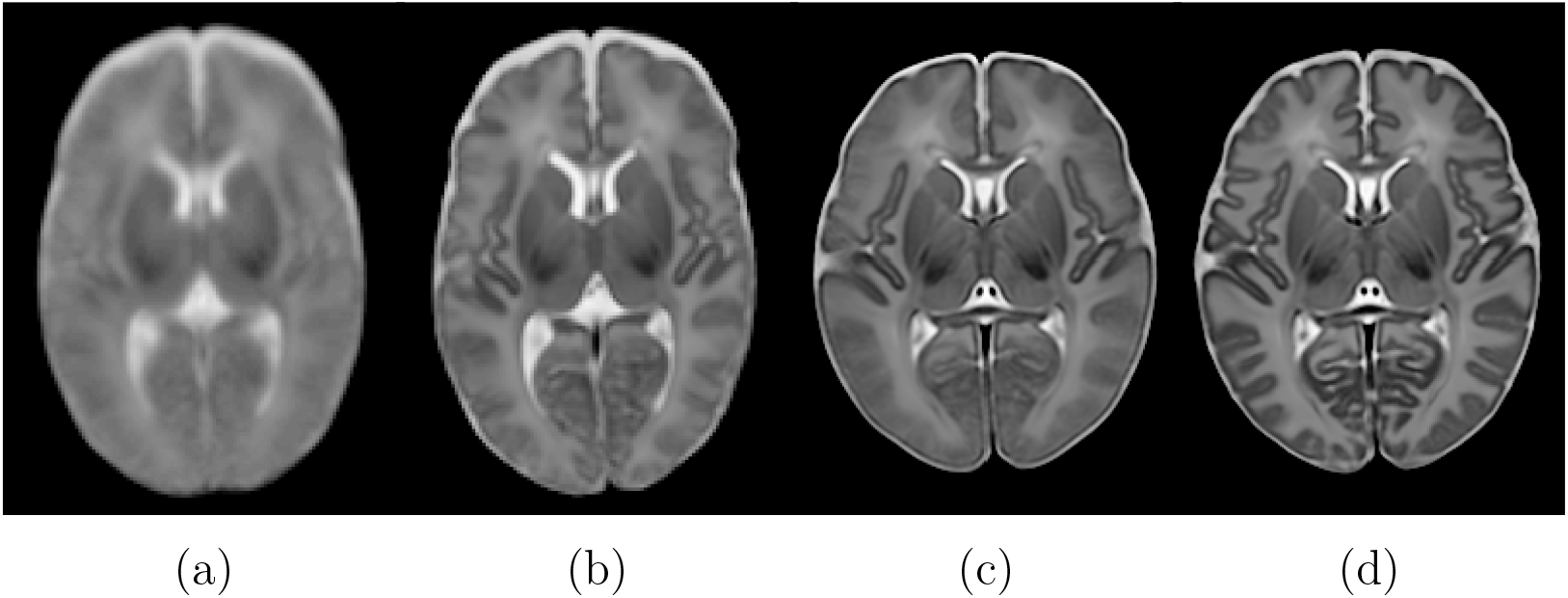
Qualitative comparison of mean T2w image at 40 weeks of (a) atlas created by Kuklisova-Murgasova et al. (2011) from 142 preterm-born neonates using Log-Euclidean mean of affine transformations with constant kernel width of 2 weeks, (b) atlas created by Serag et al. (2012a) from 204 preterm-born neonates using arithmetic mean of pairwise FFDs with adaptive kernel widths, (c) our pairwise atlas constructed from 275 term-born neonates using the Log-Euclidean mean of SVFFDs also with adaptive kernel widths, and (d) the atlas constructed from the same brain images using our groupwise approach.

## 4. Discussion

In this work, we have proposed a new iterative method for the construction of a spatio-temporal atlas of neonatal brain morphology with markedly increased anatomical detail when compared to neonatal brain atlases that were made publicly available before^2^. This new approach is compared to our previously proposed techniques based on mean deformations derived from weighted spatial mappings between all unique pairs of acquired brain images (Serag et al., 2012a; Schuh et al., 2014). Moreover, similar to the fetal atlas construction of Gholipour et al. (2017), which extended the single-time point SyGN approach to a spatio-temporal model using the adaptive temporal kernel regression used by Serag et al. (2012a), we show that an individual refinement of discrete time points, by means of registering the brain images to each respective template brain image separately, may introduce implausible temporal inconsistencies. The atlas constructed with our groupwise technique does not suffer from these temporal inconsistencies.

For the comparison of the iterative approaches, we used our SVFFD algorithm instead of the greedy SyN method utilised by Gholipour et al. (2017) for several reasons. First, our main goal was to demonstrate the temporal inconsistencies that may be introduced by independent non-linear registrations to different time points due to the highly non-convex nature of the energy functions. Secondly, although SyN (Avants et al., 2008) has been demonstrated to be among the best performing deformable registration methods (Klein et al., 2009; Ou et al., 2014), the greedy optimisation can only provide us with the final displacement field and its inverse, instead of a (non-stationary) velocity field, unless the intermediate instantaneous velocity fields were to be recorded. A mean diffeomorphism corresponding to the average velocity fields can thus not be computed. The properties of a velocity-based parametrisation are hereby traded in for a more efficient registration, which does not require the costly space-time optimisation of LDDMM (Beg et al., 2005). A further drawback of a greedy optimisation, even when the intermediate update fields are preserved, is that it may lead to a suboptimal flow of diffeomorphisms in terms of the deviation from the shortest path connecting the identity mapping from the final deformation along the manifold of diffeomorphisms. When defining a mean deformation based on these velocity fields, a deformation corresponding to a longer path contributes more to the mean shape deformation than a flow that follows the geodesic. Finally, although SyN is a greedy method, it has an average runtime of several hours^3^. For comparison, our SVFFD algorithm has an average multi-threaded runtime of 8-15 min when executed on a machine with 8 CPU cores, and the result is a SVF based on which a mean diffeomorphism can be computed.

It is worthwhile to note that in order to increase the sharpness of the constructed mean images, the Laplacian sharpening implemented in the Insight Segmentation and Registration Toolkit (ITK)^4^ (Yoo, 2004) is applied. We noticed that this filter is used by the SyGN implementation in ANTs, on which the results presented by Gholipour et al. (2017) are based, and therefore adopted this post-processing step. Note also that the same edge sharpening has been done for all atlases compared in this work, except for the publicly available atlas templates shown in figures 16a and 16b.

## 5. Conclusions

We have proposed two approaches for the construction of a spatio-temporal atlas of the developing human brain based on the unbiased SVFFD algorithm. Our first approach, presented in Schuh et al. (2014), improved upon an earlier method developed by Serag et al. (2012a). Unlike the initial technique using the FFD algorithm (Rueckert et al., 1999) implemented in IRTK and arithmetic mean of displacements (Seghers et al., 2004), we utilised the SVFFD algorithm and Log-Euclidean mean of SVFs (Arsigny et al., 2006).

This preserves the topology of the brain anatomies and ensures inverse consistency of the computed spatial mappings. Using MIRTK for both pairwise methods, we demonstrated slight improvements in cortical entropy measures using the SVFFD algorithm. A disadvantage of these approaches is, however, the quadratic computational complexity with respect to the number of brain images. A conceptual limitation is the reliance on accurate pairwise correspondences. We thus proposed a new groupwise method, which demonstrates better temporal consistency when compared to other spatio-temporal extensions of iterative methods (Gholipour et al., 2017), and results in average brain shape and intensity images with significantly more cortical details.

We conclude with a remark by Sled and Nossin-Manor (2013), that an inherent difficulty in brain atlas formation is the normal variability among individuals, and the uncertainty of the PMA. In time, these are mitigated by larger datasets acquired shortly after birth. The acquisition and public dissemination of more than a thousand high-quality brain MR images of the developing brain from 20 to 44 weeks PMA is the goal of the dHCP. The method presented in this work facilitates the construction of a spatiotemporal atlas of brain morphology from this and other large datasets.

An open source implementation of the SVFFD algorithm and our atlas construction methods are publicly available as part of MIRTK^5^. The atlases constructed from the dHCP cohort will be publicly available for download from https://data.developingconnectome.org.

## Acknowledgements

This work has received funding from the European Research Council under the European Union’s Seventh Framework Programme (FP/2007-2013) [grant number 319456]. We are grateful to the families who generously supported this trial. The work was supported by the NIHR Biomedical Research Centers at Guy’s and St Thomas’ NHS Trust.

## Appendix A. Cubic B-spline parametrisation of SVF

The SVF, whose parameters are optimised by the SVFFD algorithm, is

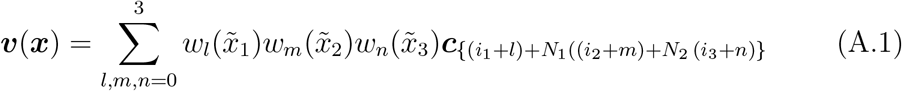

with coefficient vectors 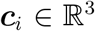. The local coordinates of 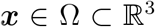 with respect to the control point lattice of size *N*_1_*N*_2_*N*_3_ are given by a rotation ***R*** ∈ 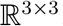, translation 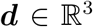, and anisotropic scaling by factors corresponding to the uniform spacing δ_d_ of control points along the *d*-th axis of the lattice, i.e., 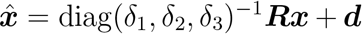. The coordinates of *x* relative to the edges of the cell of the lattice that it is contained in are then given by 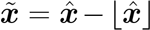 and 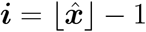 is the multi-index of the “leftmost” control point with nonzero weight. The weights 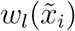 are defined in terms of the polynomials *B_i_* of the cubic B-spline function with knots at −2, −1, 0, 1, and 2. These are derived using Cox-de Boor’s recursion formula (de Boor, 2001), i.e.,

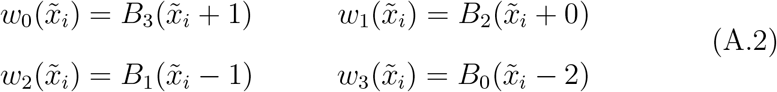

where

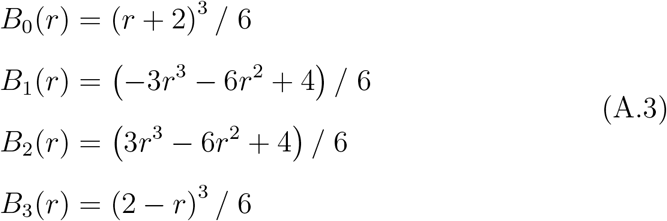

## Appendix B. Iterative pairwise affine registration

The affine transformation 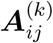 is obtained by registering the *j*-th to the *i*-th image after application of the previous mean transformations, i.e., by registering the images 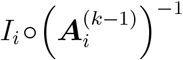 and 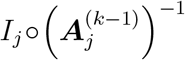. The blurring effect caused by interpolation, when resampling the images in the reference space, is avoided through composition with the transformations 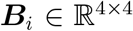, which map each image sampling space to the common world space. Specifically, the map relating a sampling point 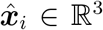 in the space of the i-th image, with axes parallel to the sampling grid and scaled according to the image resolution, to its corresponding point 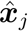 in the *j*-th image is given by

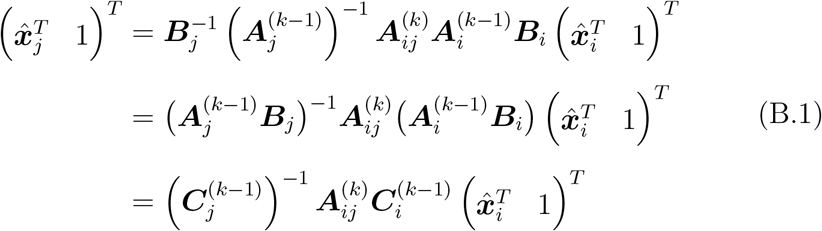

Registering images with pre-transformations 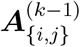 is thus equivalent to registering images with the composite image to world maps denoted by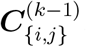. The registrations at the *k*-th iteration find the affine mappings 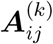 that maximise the NMI of each image pair.

## Footnotes

1 https://github.com/BioMedIA/IRTK (last accessed: Oct 13, 2017)

2 http://brain-development.org/brain-atlases/

3 An average runtime of ten hours is reported in Tustison and Avants (2013).

4 https://itk.org/

5 https://biomedia.doc.ic.ac.uk/software/mirtk/

